# Fast, high-throughput production of improved rabies viral vectors for specific, efficient and versatile transsynaptic retrograde labeling

**DOI:** 10.1101/2021.12.23.474014

**Authors:** Anton Sumser, Maximilian Joesch, Peter Jonas, Yoav Ben-Simon

## Abstract

To understand the function of neuronal circuits it is crucial to disentangle the connectivity patterns within the network. However, most tools currently used to explore connectivity have low-throughput, low-selectivity, or limited accessibility. Here, we report the development of an improved packaging system for the production of the highly neurotrophic RVdG_envA_-CVS-N2c rabies viral vectors, yielding titers orders of magnitude higher with no background contamination, at a fraction of the production time, while preserving the efficiency of transsynaptic labeling. Along with the production pipeline, we developed suites of “starter” AAV and bicistronic RVdG-CVS-N2c vectors, enabling retrograde labeling from a wide range of neuronal populations, tailored for diverse experimental requirements. We demonstrate the power and flexibility of the new system by uncovering hidden local and distal inhibitory connections in the hippocampal formation and by imaging the functional properties of a cortical microcircuit across weeks. Our novel production pipeline provides a convenient approach to generate new rabies vectors, while our toolkit flexibly and efficiently expands the current capacity to label, manipulate and image the neuronal activity of interconnected neuronal circuits *in vitro* and *in vivo*.

---

Addressing the complexity and underlying structure of neuronal networks remains one of the biggest challenges in modern neuroscience, as this knowledge is essential for the understanding of circuit functionality in health and disease^1, 2^. Short- and long-range connectivity between populations of neurons in the central and peripheral nervous systems, can be mapped with numerous existing techniques, with varying degrees of simplicity, efficiency and reliability^3, 4^. While recent advances in viral technology, such as anterograde and retrograde AAVs, now enable genetic targeting of populations based on their projection pattern^5, 6^, they can only be used for analysis of macrocircuits, due to their lack of specificity. In contrast, monosynaptic tracing technologies, based on engineered rabies virus, enable cell-type specific labeling of presynaptic partners, making it one of the most powerful toolkits to genetically dissect neuronal populations. This process is achieved in two steps: in the first, the avian TVA receptor and the rabies glycoprotein (G) are co-expressed in a predesignated population of cells, usually via administration of an AAV vector (Fig. 1a) and in the second step, G-deleted rabies viral vectors, pseudotyped with the Rous Sarcoma Virus’s envelope glycoprotein A (envA) are introduced to the region with TVA and G expressing neurons (Fig. 1b), resulting in propagation of rabies particles exclusively from these starter cells to their presynaptic partners, but not to disynaptically connected neurons, and rarely to post-synaptic targets^7^, regardless of their physical proximity^8, 9^ (Fig. 1c).

**Figure 1:**
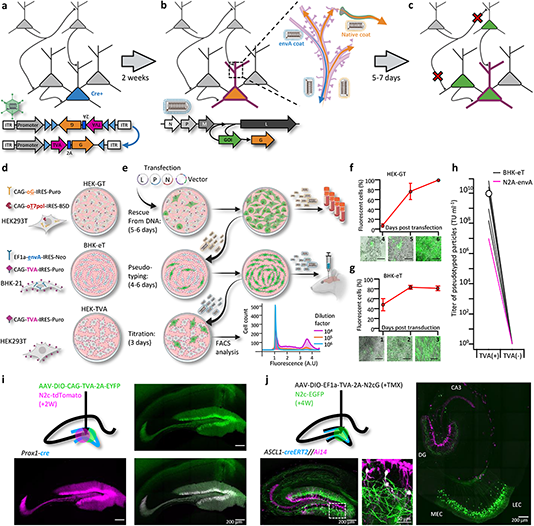
A new viral packaging system for fast, clean and high-throughput production of RVdG-CVS-N2c vectors. **a-c,** A schematic representation of the experimental workflow for achieving cell type-specific trans-synaptic retrograde labeling using G-deleted, envA-pseudotyped rabies viral vectors: Genetic dissection of cre^+^ neurons (blue) for conditional expression of TVA and G (a); targeting of RVdG_envA_ particles to labeled neurons for expression of a gene of interest (GOI, b) and subsequent propagation of native-coat RVdG particles from starter cells to their presynaptic partners (c). **d,** A schematic representation of the three different cell lines designed for rescue (HEK-GT), pseudotyping and amplification (BHK-eT) and titration (HEK-TVA) of RVdG viral particles, alongside the transgenes used to generate them. **e,** Schematic representation of the production process and timeline. L,P and N represent the plasmids encoding the corresponding rabies genes and V represents the vector plasmid. **f,** Quantification of the time course for the rescue stage, starting at day 4 after transfection of viral plasmids. **g,** Quantification of the time course for amplification of pseudotyped particles, starting at day 1 after transduction with native-coat particles. **h,** Quantification of the average titer of concentrated pseudotyped stock from 19 individual productions, with comparison to the titer of a representative production of RVdg-CVS-N2c virus, produced using N2a cells (magenta). Lines represent titers of individual productions. **i,** Schematic of the injection scheme (top left) and corresponding representative confocal images demonstrating efficiency and specificity of CVS-N2c delivery to DGCs. **j,** Schematic representation of the injection scheme (top left) and corresponding representative confocal images demonstrating efficiency and specificity of CVS-N2c-mediated retrograde labeling from adult-born DGCs.

While this approach ostensibly identifies presynaptic partners associated with a predesignated starter population, inefficiencies in the widely used SAD-B19 strain, along with potential transmission biases across cell types^10^ suggest the traced cells might represent only a fraction of the complete presynaptic population, rendering smaller, more distributed projections more difficult to identify. Furthermore, since large quantities of native-coat particles are routinely used to initiate the pseudotyping step of the viral production protocol^11^ background contamination of native-coat particles in the pseudotyped stock is nearly unavoidable, which can result in false identification of projecting populations directly labeled by native-coat particles and not transsynaptically through the starter cells.

Recently, the deployment of the highly neurotropic CVS-N2c rabies strain for monosynaptic tracing was shown to identify 5 to 20 fold more presynaptic cells per starter cell than SAD-B19, enabling a more comprehensive analysis of the diversity of presynaptic cells innervating a predesignated starter population. In addition, this strain was shown to be less neurotoxic, making it more suitable for experiments in behaving animals. However, the production process for these vectors is time-consuming and yields low viral titers^12^, presumably due to the use of neuronal cell lines for the various amplification and packaging steps. This limitation has so far restricted more widespread use of the superior CVS-N2c vector in circuit mapping experiments, particularly in its more useful pseudotyped form, whose preparation is even lengthier and the titers lower still.

Here, we report the development of a new packaging system, allowing expedited production of high-titer RVdG_envA_-CVS-N2c particles, free of background contamination from native-coat particles, and show that these vectors retain their superior expansion efficacy when compared to SAD-B19 vectors. We also report an extended toolkit of AAV and CVS-N2c vectors which can be applied to a wide range of experimental paradigms, and demonstrate their efficacy in uncovering hidden neuronal connections, even in heavily explored circuits.

## Results

### Improved packaging for RVdG viral vectors

Prolonged and cumbersome production has been a major limitation for G-deleted rabies virus vectors and most profoundly so for the CVS-N2c strain. Because CVS-N2c vectors exhibit enhanced neurotropism and retrograde labeling over SAD-B19, improvements in the speed and quality of the various N2c production stages are necessary to realize the full potential of rabies viral vectors for mapping synaptic circuits. Here we introduce a production protocol, based on two new packaging cell lines we have developed: 1) “HEK293-GT” cells, based on HEK293T cells stably expressing the SAD-B19 optimized glycoprotein (oG) along with the optimized T7 RNA polymerase (oT7pol), used for the initial rescue and amplification of native-coat particles, and 2) “BHK-eT” cells, based on BHK21 cells stably expressing the envA glycoprotein, along with the TVA receptor, used for simultaneous vector pseudotyping and amplification (Fig. 1d,e). Both cell lines make use of antibiotic resistance genes and strong constitutive promoters^13^ to ensure cultures consist of purely transgene-expressing cells and that expression levels in those cells is high (Extended Data Fig. 1a-c). Furthermore, unlike previous packaging systems, the co-expression of TVA and envA in the BHK-eT cell line allows pseudotyped particles to propagate within the culture, similar to the native-coat ones (Fig. 1f,g), enabling amplification and pseudotyping of vectors in a single short step. In addition, since a very small amount of native-coat, or even trace amounts of pseudotyped virus, are sufficient in order to initiate the pseudotyping process, contamination from native-coat particles in the final pseudotyped stock can be minimized, while the titer of the pseudotyped stock can be exceedingly high (Fig. 1h). Together, these new cell lines enable significantly faster production times, higher titers and less background contamination than both other currently used methods for production of either CVS-N2c or SAD-B19 (Supplementary Table 1).

To test the efficacy and transduction exclusivity of these new vectors, we first expressed a cre-dependent TVA-2A-EYFP cassette, which lacks the rabies glycoprotein, in dentate granule cells (DGCs), using prox1-cre mice^14^. Subsequent delivery of high-volume (0.5 µl), high-titer (∼3X10^10^ TU/ml) RVdG_envA_-CVS-N2c-tdTomato vectors resulted in exclusive expression of tdTomato in EYFP-labeled neurons in the DG (Fig. 1i). To evaluate whether these particles efficiently propagate transsynaptically, while retaining their target specificity, we performed an additional experiment in-which a TVA-2A-N2cG cassette was expressed in a small population of adult-born DGCs, using Ascl1-creERT2 line crossed with the Ai14 tdTomato cre-reporter line^15^. Four weeks after the Adeno-Associated Viral (AAV) vectors were delivered to the DG, along with a single i.p. injection of tamoxifen (TMX) to induce recombination in the CreERT2 line, RVdG_envA_-CVS-N2c-EGFP vectors were introduced to the same location. This manipulation resulted in both highly specific targeting of tdTomato^+^ adult-born DGCs in the inner cell layer, as well as robust retrograde labeling of afferents to these starter cells. This high efficacy manifested as widespread and specific labeling of layer-II neurons of the medial and lateral entorhinal cortices (MEC and LEC respectively), hilar mossy cells, CA3 pyramidal cells and a number of DG-projecting subcortical regions, including the medial septal (MS), the supramamilary (SuM) and the raphe (RN) nuclei (Fig. 1j and Extended Data Fig. 2a,b), largely consistent with several previous reports^16, 17^. These experiments provide evidence that the new packaging system we designed is capable of rapidly producing high-titer and high-quality RVdG_envA_ vectors for transsynaptic retrograde neuronal labeling.

### Comparative retrograde labeling efficiency of RVdG-CVS-N2c particles

RVdG-CVS-N2c vectors were originally produced in the neural progenitor cell line Neuro2A (N2A), under the premise that this would improve neurotropism of the assembled viruses, enhancing their trans-synaptic transfer rates and reducing their neurotoxicity^12^. The new packaging cell lines we developed were designed to increase production speed and efficiency, but the possibility remains that the non-neuronal origin of the BHK-21 cells might compromise the superior properties of CVS-N2c vectors. To evaluate any differences in retrograde labeling efficiency between the vectors assembled using the two approaches, we expressed a TVA-2A-N2cG cassette in CA1 pyramidal neurons and subsequently transduced them with a cocktail of two different RVdG_envA_-CVS-N2c vectors expressing either EGFP or dTomato, in equal titers, amplified and pseudotyped using either N2A-based or the new HEK-GT/BHK-eT based packaging systems (Fig. 2a). This strategy resulted in highly efficient and specific retrograde labeling of CA1 afferents in the CA3, LEC and medial septum (MS) in equal proportions for both vectors (Fig. 2b,c,h).

**Figure 2:**
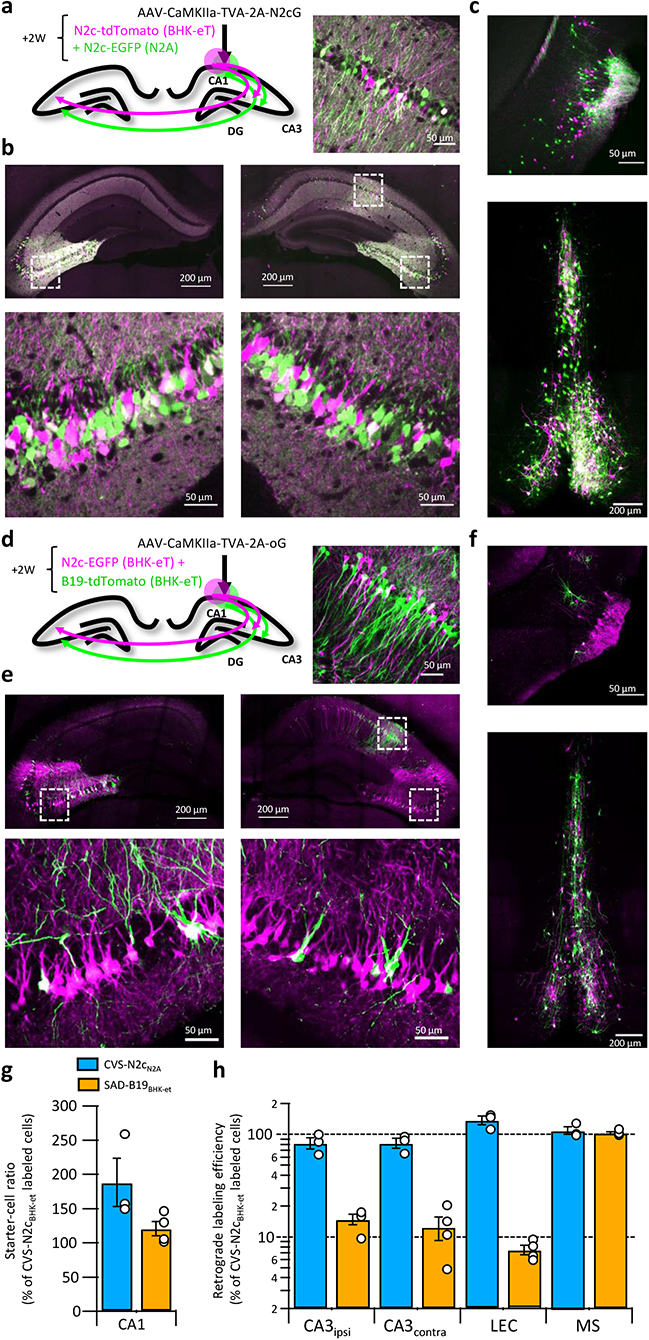
Retrograde labeling efficiency is determined by the viral strain, and not the packaging cell lines. **a,** Schematic illustration of the injection scheme, designed for comparison of retrograde labeling efficacy between CVS-N2c vectors produced using either the N2a-based or BHK-based packaging cell lines, propagating using the N2c glycoprotein. **b,** Representative confocal images of the ipsi-, and contra-lateral hippocampus. Expanded images of the CA3 region of both hemispheres and the CA1 of the injection site correspond to the areas delineated by white rectangles. **c,** Representative confocal images of the LEC (top) and the septal complex (bottom). **d-f,** Same as (a-c) but for comparison of CVS-N2c and SAD-B19 vectors, both produced with BHK-based packaging cell line and propagating using the SAD-B19 optimized glycoprotein (oG). **g,** Summary bar plot showing the ratio of 1^st^ order starter cells in the CA1 pyramidal layer, between neurons labeled with either CVS-N2c_N2a_ (cyan) or SAD-B19_BHK-et_ (orange) and the neurons labeled with CVS-N2c_BHK-et_. **h,** Summary bar plot showing the differences in retrograde labeling efficiency, under both injections schemes described in (a) and (d). All values were normalized to the ratio of starter cells shown in (g). N= 4 and 3 animals for the N2c-N2c and N2c-B19 comparisons, respectively. Data shown as mean and SEM with white circles denoting individual animals.

To confirm that these CVS-N2c vectors remain more efficient than the widely-used SAD-B19 strain, as was previously shown for vectors produced in N2A cells^12, 18^, the previous experimental design was repeated, using the optimized B19 glycoprotein (oG)^19^ and RVdG_envA_-SAD-B19 vectors (Fig. 2d). Here, a strikingly different result was observed, with afferents labeled with CVS-N2c visibly outnumbering those labeled with SAD-B19 in most regions tested (Fig. 2e,f,h). Quantification of cell numbers in these experiments shows that while the ratio of 1^st^ order CA1 neurons between the two compared vectors remained low in both experiments (Fig. 2g), the ratios of 2^nd^ order neurons in the tested sets of regions differed substantially, with only minor differences observed when the two CVS-N2c vectors were compared, but differences close to an order of magnitude observed when CVS-N2c and SAD-B19 vectors were compared (Ipsilateral CA3: 14.43 ± 1.67 %; Contralateral CA3: 11.95 ± 2.85 %; LEC: 8.17 ± 0.85 %; MS: 100.88 ± 4.54 %; Fig. 2h). A notable exception is seen in the projection from the MS, in which ratios between the SAD-B19 and N2c remained identical. This effect could be the result of differences in structure of the synaptic contacts between MS and CA1 neurons or in the projection’s connectivity scheme. Since we observed that CVS-N2c vectors propagate less efficiently when using the oG, as opposed to their endogenous N2cG (Extended Data Fig. 3a-c), the actual differences in retrograde labeling between CVS-N2c and SAD-B19 are likely to be substantially higher, matching those previously reported^12^.

### Identification of intra-hippocampal projections to DGCs

To further test and validate the throughput and sensitivity of this tool, we performed additional retrograde labeling experiments from the DG, in order to see whether we will be able to corroborate and expand on previous reports, describing non-canonical inhibitory projections it receives from inhibitory neurons in the *Stratum Oriens* (*S.O.*) and *Stratum Lacunosum Moleculare* (*S.LM.*) of the CA1^20–23^. These findings are all based on reconstruction of the axonal plexus of biocytin-labeled neurons and, while informative, this approach has low-throughput and mostly lacking information about the identity of the postsynaptic partner. To see if we could locate and identify these cells, we targeted RVdG_envA_-CVS-N2c-tdTomato vectors to DGCs of Prox1cre mice crossed with GAD1-EGFP transgenic mice, in which GABAergic neurons are fluorescently labeled^24^, and examined the regions outside of the DG for colocalization (Fig. 3a). Consistent with these reports, we found extensive labeling of neurons with N2c-tdTomato, whose somata were located in the *S.O.* and *S.LM.,* but not in the *Stratum Pyramidal* (*S.P.*) or *Radiatum* (*S.R.*) layers. In addition, we uncovered a third population of DG-projecting neurons in the superficial most layer of the subiculum (Fig. 3b). Analysis of colocalization has revealed that while half of all retrogradely labeled neurons found in the *S.O* and *S.LM.* were also positive for EGFP, in the subicular population, which accounted for more than a third of all intrahippocampal projecting cells, colocalization was almost completely absent (Fig. 3c), suggesting that this population consists of excitatory neurons. AAV vectors expressing EGFP under either the inhibitory mDLX or the excitatory CaMKIIa promoters, injected into the superficial CA1 or superficial subiculum, respectively, confirmed the existence of axonal branching in the DG (Extended Data Fig. 4a,b). Identification of these populations following retrograde labeling from adult-born DGCs ventrally located DGCs (Extended Data Fig. 4c,d) further support the existence of these connections. Taking advantage of the high-throughput nature of our labeling approach, we immunostained sections from labeled animals for two of the most prominent interneuron markers, Parvalbumin (Pvalb) and Somatostatin (Sst) and found that while half of *S.O.* retrogradely-labeled neurons were positive for Sst, none of the labeled cells in both *S.O.* and *S.LM* colocalized with Pvalb (Fig. 3d,e). This is consistent with the known projection of *S.O.* Sst neurons to the *S.LM.*, which borders on the dentate gyrus, while axons of Pvalb neurons in the CA1 project mainly to the pyramidal cell layer and have few, if any axonal arborization in the *S.LM.*^25^ and are therefore much less likely to branch into the DG.

**Figure 3:**
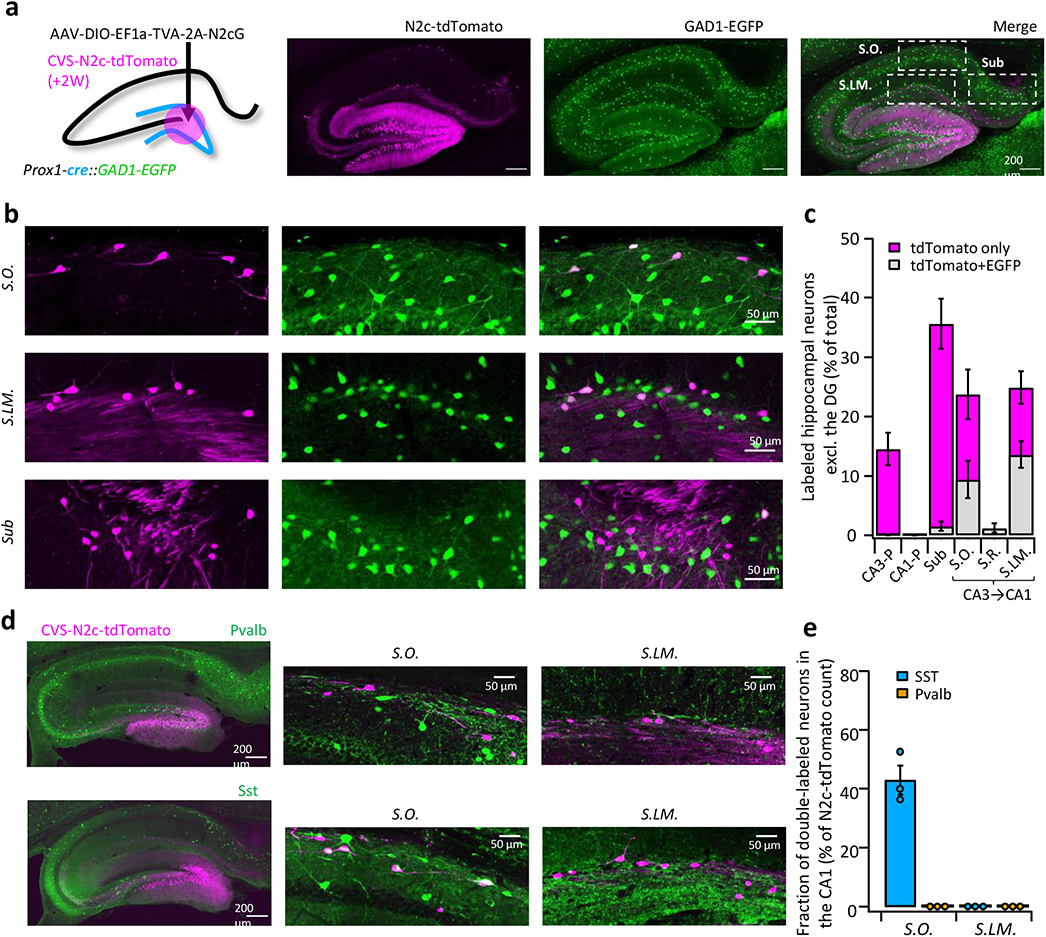
High-throughput retrograde labeling with CVS-N2c enables identification of non-canonical projections to DGCs. **a,** Schematic illustration and representative confocal images, describing the injection scheme designed to target DGCs for retrograde labeling in an interneuron reporter line. **b,** Representative confocal images (left) of the regions highlighted in (a) showing retrogradely-labeled neurons along specific hippocampal layers and their overlay with the interneuron-specific marker. **c,** Summary bar plot showing the distribution of DG-projecting hippocampal neurons outside of the DG (magenta) and of them, the fraction of double-labeled neurons (grey). Calculation of cell numbers in the dendritic cell layers combined cells along the entire proximo-distal hippocampal axis, from CA3 to CA1. N = 189 cells from 3 animals**. d,** Representative parasagittal sections of the hippocampus following retrograde labeling from the DG with CVS-N2c-tdTomato, along with immunolabeling of parvalbumin (Pvalb, top), Somatostatin (Sst, bottom). Expanded view of the *S.O.* and *S.LM.* are shown to the right of each image. **e**, Summary plot describing the proportion of Pvalb or Sst positive neurons of the total CVS-N2c labeled neurons in the *S.O.* or *S.LM.* of the CA1. N = 125 cells from 3 animals.

### Differential targeting of neuronal populations for retrograde labeling

The abundance of population-specific cre/flp driver mouse lines allows for a wide range of possible labeling experiments. However, often such lines are either unavailable, or insufficiently specific for the experimental requirements. To allow for a broader use of this tool, we have developed additional AAV vectors driving expression of the TVA-2A-N2cG cassette which are capable of achieving greater specificity.

First, we wanted to be able to compare between two genetically distinct, yet spatially overlapping populations. To this aim, we designed AAV vectors which contain a cre-off mechanism, using a single-floxed, excisable open reading frame (SEO) to drive transgene expression under control of the excitatory neuronal CaMKIIa promoter (Extended Data Fig. 5a). We tested this tool on CA1 pyramidal neurons of Calb1-cre mice, in which Cre-Recombinase is expressed exclusively in deep, but not in superficial CA1 neurons^26, 27^ (Fig. 4a). Parallel retrograde labeling from these two subpopulations of CA1 neurons revealed that while both receive relatively equal input from CA3 neurons, deep CA1 neurons receive substantially greater input from LEC-III neurons (Fig. 4b), confirming previous findings obtained using lower-throughput approaches^26, 28^.

**Figure 4:**
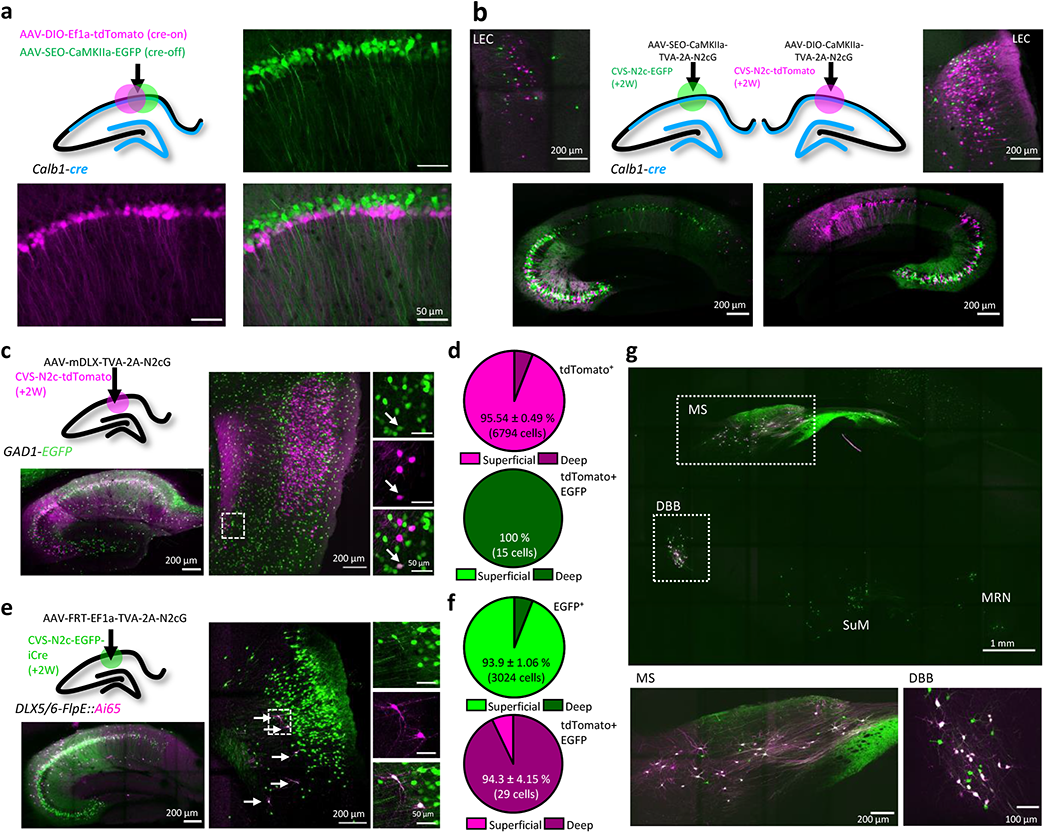
An AAV vector suite for targeting multiple and diverse neuronal populations. **a,** Graphical representation (top left) and representative confocal images, demonstrating differential targeting of superficial and deep CA1 pyramidal neurons using a combination of cre-on and cre-off AAV vectors. **b,** Graphical representation (top center) for dual retrograde labeling from superficial (bottom left, green) and deep (bottom right, magenta) CA1 pyramidal neurons and the resulting distribution pattern of their corresponding projection neurons in the EC (top left and top right). **c,** Graphical representation of the viral injection scheme for mapping inputs into hippocampal inhibitory neurons (top left), and representative parasagittal images of labeled cells in the HC (Bottom left) and LEC (right). Expanded images show a double-labeled neuron in EC-VI. **d,** Distribution of all retrogradely labeled cells (top) and of double-labeled cells only (bottom) among the superficial layers II and II and the deep layers V and VI of the LEC. N = 8 sections/3 animals. **e,** Same as (c). **f,** Same as (d) for the experiments described in (e). N = 8 sections/3 animals. **g**, A representative parasagittal image of deep brain structures following the injection scheme described in (e). SuM – Supramammilary Nocleus; MRN – Median Raphe Nucleus.

Next, we wanted to test whether our vectors could also label projections to inhibitory neurons. To attain interneuron-specific retrograde labeling, we first used AAV vectors to express the TVA-2A-N2cG cassette under control of the interneuron-specific mDLX promoter^29^ in the CA1 of GAD1-EGFP mice. A subsequent injection of CVS-N2c-tdTomato revealed widespread labeling of 2^nd^ order neurons both within the hippocampal CA1 and CA3 fields, as well as in the EC (Fig. 4c and Extended Data Fig. 5b). An additional analysis of co-labeling in the EC revealed a small population of long-range inhibitory projection neurons, located preferentially in the deeper layers Va and VI (Fig. 4d), whose existence has previously been reported^30, 31^ but their location within the EC has so far remained unknown. In order to cross-validate and expand on these findings, we crossed Dlx5/6-flpE mice with the double cre+flp, tdTomato reporter line Ai65^32^. Hereby, using injections of AAV vectors with flp-dependent TVA-2A-N2cG expression cassette, followed by RVdG_envA_-CVS-N2c-EGFP-iCre, we were able to highlight and isolate the inhibitory projections to hippocampal inhibitory neurons, as the double cre+flp recombination required for tdTomato expression can only take place in inhibitory neurons transduced by the rabies virus (Fig. 4e). In line with our previous findings, we show that while the majority of excitatory cortical input to hippocampal inhibitory neurons originated in the superficial layers II and III, long-range inhibitory projection neurons are almost exclusively found in the deeper layers Va and VI (Fig. 4f). Further examination of labeling patterns of 2^nd^ order neurons in deep-brain regions showed that in the MS and DBB, a large fraction of cells are double-labeled, confirming that our labeling strategy for isolation of long-range inhibitory projections is exhaustive (Fig. 4g). This strongly suggests that while the deeper cortical layers give rise to both inhibitory and excitatory hippocampal projections, whereas the excitatory projection originated from a substantially larger population. This is particularly unexpected, as to-date, only spurious evidence existed for the presence of cortico-hippocampal projections, excitatory or inhibitory, arising from the deeper layers of the EC^33^.

### Bicistronic CVS-N2c vectors for efficient dual labeling

A previous report has shown that the B19 N-P linker sequence, which allows the virus to effectively separate these proteins, can also be used for separation of exogenous genes^34^. While this approach could promote an expansion of the rabies toolkit to accommodate for more complex experimental designs, it remains to be shown to whether efficient separation indeed takes place and whether, unlike the use of 2A peptides or the IRES sequence, expression levels of the individual proteins remain unaltered. To test the feasibility and efficacy of this approach, we designed new CVS-N2c bicistronic plasmids, to drive co-expression of a nuclear-localized EGFP (nl.EGFP) with either tdTomato (Fig. 6a) or a synaptophysin-tethered EGFP (SypEGFP, Fig. 6b) separated by the CVS-N2c N-P liker sequence. Specific delivery of these vectors to hippocampal DGCs using targeted AAV expression of the cre-dependent TVA-2A-N2cG cassette in Prox1-cre mice, revealed that in both cases, the individual proteins were efficiently expressed in a compartment-specific manner (Fig. 5a,b and Extended Data Fig. 6a-c).

**Figure 5:**
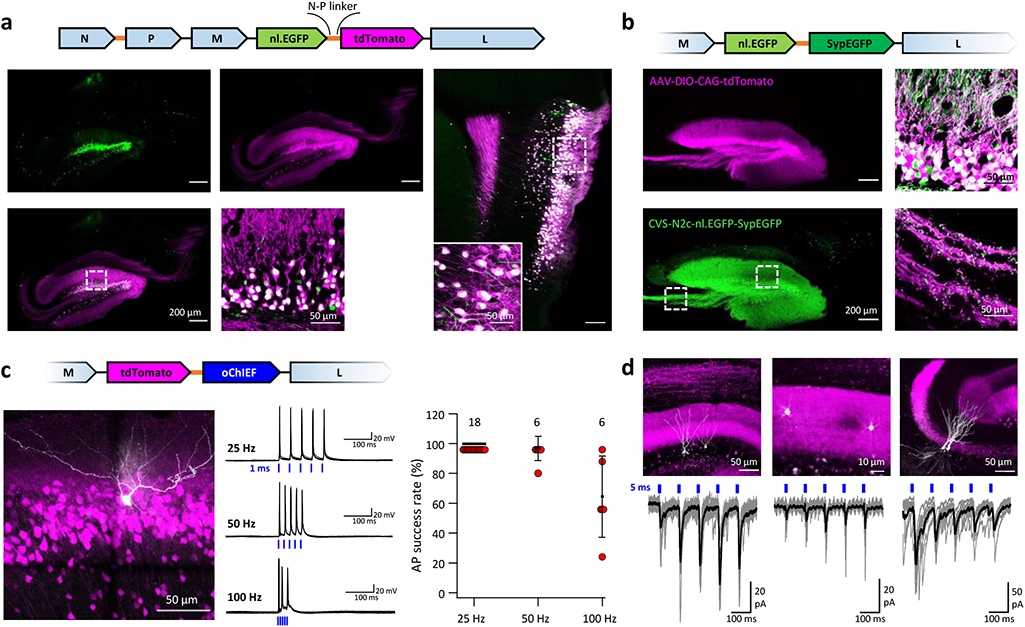
An extended suite of RVdG-CVS-N2c vectors for bicistronic expression of fluorescent markers and optogenetic effectors. **a,** Schematic illustration of the vector sequence, designed to drive independent bicistronic expression of a nuclear-localized EGFP (nl.EGFP) alongside tdTomato, using the N-P linker sequence (top). Representative confocal images of the HC (bottom right) and EC (bottom left) following retrograde labeling from the DG, demonstrate the differential localization, indicating effective separation of the fluorophores. **b,** Schematic diagram of a bicistronic nl.EGFP + SypGFP CVS-N2c vector (top) used for retrograde labeling from the DG (right panels) and representative confocal images demonstrating dual nuclear and synaptic localization of EGFP in the dentate granular and molecular layer (top right image) and purely synaptic localization at the mossy fibers terminals (bottom right image). **c,** Schematic diagram of a bicistronic tdTomato + oChIEF CVS-N2c vector (top) used for retrograde labeling from the DG, and a representative image of a biocytin-filled neuron (white) in MEC-II (bottom right). Representative traces from 10 overlaid recordings at different frequencies (bottom center) and summary plot (bottom left) of the action potential failure rate for recordings made 6-7 days after introduction of RVdG, demonstrate the light responsiveness of transduced cells. **d,** Representative confocal images (top) of DGCs (left), DG molecular layer interneurons (center) and CA3 pyramidal neurons (right) and their synaptic responses to optogenetic activation of the perforant path (bottom) following retrograde labeling from the dorsal DG with the bicistronic CVS-N2c-tdTomato-oChIEF vector.

**Figure 6:**
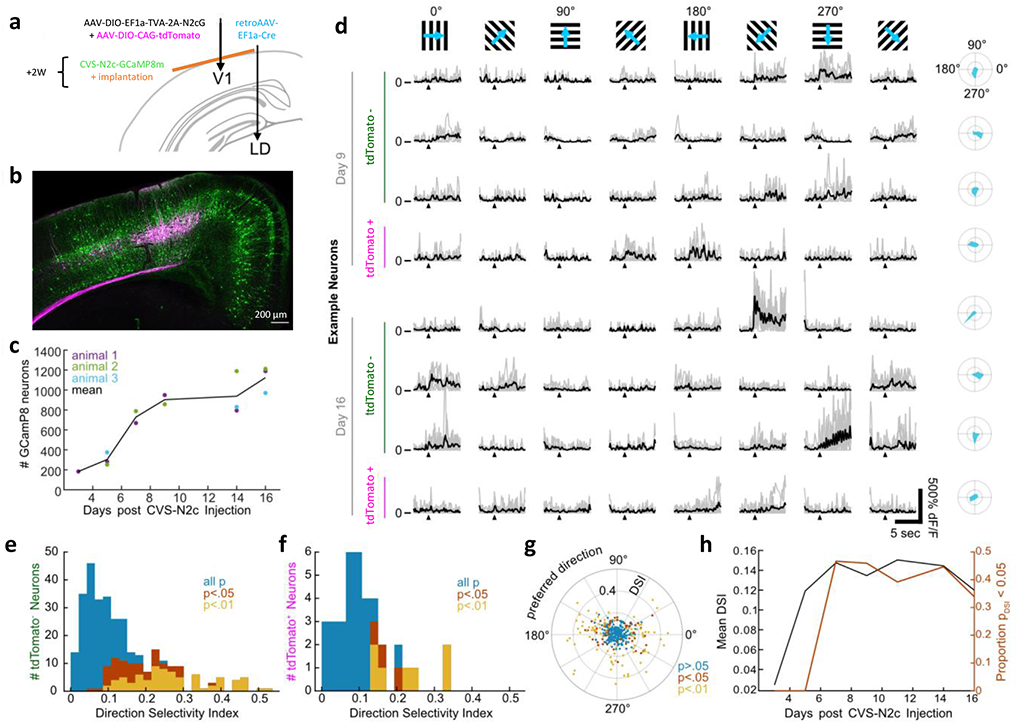
New CVS-N2c vectors for *in vivo* two-photon long-term imaging of activity in a cortical microcircuit. **a,** Illustration of the injection scheme for labeling projections onto the V1 region’s layer V neurons with GCaMP8m. **b,** A representative confocal image of a coronal section following the injection scheme described in (a). **c,** GCaMP8m positive neuron numbers over recording days. **d,** In vivo two-photon imaging in presynaptic neurons. Drifting gratings in 8 directions (top row). Trial responses (grey) and average (black) for example neurons on day 9 and day 16, stimulation starts at black triangle. Polar plot of the directional responses in right column. **e,** Histogram of direction selectivity for tdTomato negative (2^nd^ order) neurons. **f,** Same as in E for tdTomato positive (starter) neurons. **g,** Polar scatter plot of direction selectivity (radial) over preferred direction (angular) for all recorded neurons. **h,** Mean DSI and proportion of significant direction selectivity over recording days.

Next, we capitalized on this result to create new bicistronic vectors for dual expression of the optogenetic actuator ChIEF^35^, along with a fluorescent protein, speculating that the increased expression levels and the untethering of the fluorophore will lead to greater responsivity of labeled neurons to optogenetic stimulation, as well as facilitate their identification. We again targeted DGCs for retrograde labeling using these new vectors and recorded the action potential success rate of labeled neurons in the EC, following five light pulses, each 1 ms long, at varying frequencies. As predicted, action potentials could be reliably generated at much higher stimulation frequencies and at a much earlier point following stimulation onset, than has previously been shown for both B19 and N2c vectors^12, 34^ (Fig. 5c). By recording synaptic responses to optogenetic stimulation from several neuronal population sharing the same input as DGCs, such as neighboring, non-transduced DGCs, CA3 pyramidal neurons and dentate gyrus molecular layer interneurons, we also show that this tool can be reliably used for exploring circuit motifs, and also potentially for effective circuit-based manipulation in behaving animals (Fig. 5d).

### *In vivo* measurements of a defined cortical microcircuit

While RVdG viral vectors are highly effective in describing circuit architecture, their neurotoxicity limits their use for many applications, in which long-term monitoring or manipulation of labeled circuit is required^4^ and while a new technology for production of non-neurotoxic RVdG vectors was recently been presented^36^, the current inability to produce these vectors in a pseudotyped form, excludes their use for cell type-specific tracing experiments. RVdG-CVS-N2c vectors have previously been shown to be less neurotoxic than the SAD-B19 strain, and compatible for prolonged imaging of neuronal activity *in vivo*^12^. However, since neurotoxicity has not been completely eliminated, it is still possible that their endogenous activity patterns become impaired within this time period, thereby impinging on results.

To address this question, we dissected a microcircuit within the primary visual cortex (V1) for *in vivo* calcium imaging, by first injecting retrogradely transported Cre-expressing AAV vectors^5^ in the laterodorsal nucleus of the thalamus (LD) and AAV-DIO-CAG-tdTomato + AAV-DIO-EF1a-TVA-2A-N2cG into the V1. A subsequent injection of RVdG_envA_-CVS-N2c-GCaMP8m into V1 resulted in specific retrograde labeling from layer V neurons in the V1 (Fig. 6a,b and Extended Data Fig. 7a,b). In the following weeks we observed the time course of GCaMP8m expression and recorded calcium transients of the labeled 1^st^ and 2^nd^ order neurons using two-photon imaging, in response to visual cues in three awake behaving mice,. While we detected the first GCaMP8m labeled neurons already at day 3, substantial labeling at approximately half-maximal numbers in all animals started 7 days after RVdG_envA_-CVS-N2c-GCaMP8m injection. Subsequently, the increase of neuronal numbers slowed and remained high until day 16 (on average 1100 neurons), when the experiments were terminated (Fig. 6c). While the most superficial tdTomato-labeled starter neurons were detectable *in vivo*, their depth did not allow a robust estimate of their total numbers in our configuration.

Each session, we recorded fluorescence time series for a subset of the GCaMP8m labeled neurons and determined their Calcium activity while presenting drifting grating stimuli to the mouse in a visual dome setup (Fig. 6d). Both tdTomato positive (putative 1^st^ order) neurons as well as tdTomato negative 2^nd^ order neurons showed similar tuning properties (Fig. 6e,f) with preferred directions slightly biased in the horizontal plane (Fig. 6g). The proportion of significantly direction tuned neurons (Fig. 6h) and signal-to-noise ratio (Extended Data Fig. 7c) remained constant from day 7 to the conclusion of the experiments at day 16. These experiments demonstrate that RVdG-CVS-N2c viral vectors can be reliably used for *in vivo* experiments in behaving animals for circuit-specific recording and manipulation of activity and delineate a time period, during which the neurons retain their normal physiological properties.

## Discussion

### Fast and efficient production of high-titer rabies virus vectors

The published production protocol for RVdG_envA_-CVS-N2c rabies viral vectors utilizes the neural precursor N2A cell line since it was hypothesized to increase neuronal tropism, thereby increasing transsynaptic spread from the starter cells. However, since this cell line is relatively more difficult to transfect and maintain than other cell lines, this choice resulted in a lengthy and cumbersome production process, with low titer yields^12^. As this premise has not been directly tested, we surmised that the increased labeling efficiency stems from the N2c strain itself, rather than its production process, and created an alternative packaging system to optimize all the different steps of production. By co-expressing mammalian-optimized versions of the genes required for the rabies amplification process, in tandem with antibiotics-resistance genes driven by strong constitutive promoters, in highly resilient and easily maintained cell lines, our method effectively minimizes production time, while simultaneously increasing viral titers by orders of magnitude. The additional co-expression of TVA and envA in the pseudotyping cell lines reduces to a minimum the amount of native-coat virus needed to initiate the process, thereby nearly abolishing the presence of these particles in the pseudotyped stock, and vastly increasing tracing specificity. This advantage is particularly relevant when performing retrograde labeling from single cells^37–39^ where a small number of non-specifically labeled neurons can potentially account for a large fraction of the entire labeled population.

RVdG-CVS-N2c vectors have previously been shown to outperform the SAD-B19 in almost any parameter measured^12, 40^, attributed to the use of neural precursor cells for their production. We show that contrary to the premise, vectors produced using the HEK-GT/BHKeT packaging system show comparable retrograde labeling efficiency to ones produced in N2A cells and significantly higher than that of SAD-B19 particles. This attribute has allowed us to efficiently find and characterize non-canonical hippocampal connections, for which only anecdotal reports have been available. While we show that neurons transduced with CVS-N2c vectors produced using our cell lines maintain viability and physiological function for at least 16 days, since no direct comparisons of physiological properties and long-term viability were made with vectors produced in the N2A line, it remains possible that packaging these vectors in N2A cells would result in reduced neurotoxicity, when compared with vector packaged in non-neuronal cell lines.

Using these new vectors, we were able to demonstrate the ease in which non-canonical projections to a target population can be teased out. We chose to focus these efforts in the DG, as the ample existing anecdotal evidence allowed us to compare our results against previously verified data^20–23^. In line with these reports, retrograde labeling from DGCs labeled two distinct populations of inhibitory neurons, residing in the *S.O.* and *S.LM* of the CA1, which mainly contain neurons projecting to the apical dendrites of CA1 neurons. In addition to these aforementioned populations, we have also observed a third population of labeled neurons along the superficial most layer of the subiculum, directly in the path of perforant path fibers. Unlike the previous ones, these cells did not express the inhibitory fluorescent indicator for GAD1, which suggests that this might be a yet undiscovered intrahippocampal excitatory projection to the DG. Since in the GAD1-EGFP line only an estimated half of inhibitory neurons express the fluorescent marker^41^, it remains a possibility that these neurons are also inhibitory but are somehow genetically indisposed to express the marker. However, their pyramidal-like morphology, spiny dendritic arbors and putative expression of the CaMKIIa promoter, all hallmarks of excitatory neurons, render this possibility unlikely.

The successful deployment of RVdG_envA_ viral vectors is mainly limited by the ability to genetically dissect the target population. While many different mouse lines have previously been for this purpose, covering the vast majority of genetically unique populations, the robust amplification capabilities of RVdG-CVS-N2c vectors requires a high degree of specificity, in order to avoid possible off-target effects. We have developed several approaches that could further improve experimental paradigms, designed to attain results that are more specific. For example, cre-off viruses with specific promoters for AAV vectors can be used together with their cre-on equivalents in order to determine which projections are specific to a target population in a given region, and which are shared by the general population of neurons in a given brain region. In addition, intersectional genomic and viral-borne recombination could be used to dissect and highlight specific sub-populations of projection neurons. Using the vectors we designed, this toolbox can be expanded further to include other restriction approaches, such as tet-controlled elements.

Complementing these tools is a new suite of mono- and bicistronic RVdG-CVS-N2c vectors, expressing a broad range of fluorophores, recombinases, synaptic markers, optogenetic actuators and genetically-encoded calcium indicators, to enable for diverse experimental purposes. (Table 1). We demonstrate the flexibility of this approach, by expressing and imaging the only recently developed calcium indicator GCaMP8m^42^ in the presynaptic neuronal population of starter neurons, that themselves project to the thalamus. Furthermore, we show that using the CVS-N2c endogenous N-P linker, we could effectively separate at least two individual elements, and possibly more. Apart from enabling better tracking of distinct cellular compartments, we also show that the separation of the fluorophore from optogenetic actuators can increase their light responsiveness, possibly as a result of improved membrane trafficking and higher expression levels. However, some of these properties should also be attributed to the improved ChR2 variant we used, which was shown to possess faster kinetics^35^. Another benefit of this separation is better identification of labeled neurons, since the untethering of the fluorophore from membrane-bound proteins leads to its predominantly cytosolic expression.

## Conclusion

We present here a comprehensive toolkit for rapid, high-throughput production of RVdG-CVS-N2c rabies viral vectors, along with an extended multipurpose AAV and CVS-N2c vector suites. These vectors now allow extensive labelling of presynaptic projections, facilitating a more comprehensive understanding of neuronal network wiring, at greater specificity, efficiency and versatility for the experimenter. Together, this system enables a fast, simple and highly effective approach for circuit mapping in the mammalian brain.

## Materials and Methods

### Generation of stable packaging cell lines

Retro-, and lentiviral vectors were produced by transfecting HEK293 cells with one of the following vectors: pMMLV-CAG-SAD-B19_optimized_G-IRES-Puro, pMMLV-CAG-optimized_T7pol-IRES-BSD, pLenti-EF1a-envA-IRES-Neo and pMMLV-CAG-TVA-IRES-Puro, along with the compatible retro-or lenti-viral GAG-Pol and the vesicular stomatitis virus glycoprotein (VSV-G), by means of calcium-phosphate precipitation. Viruses were collected and filtered 48 hours post transfection and used to transduce low-passage HEK293T or BHK-21 cells. Three days post transduction, the cells were passaged and one of the following antibiotics were added to the medium in order to select for the cells which stably express the respective construct: Puromycin dihydrochloride (3 µg/ml, Sigma Aldrich), blasticidine S hydrochloride (15 µg/ml, Sigma Aldrich) or G418 disulfate (Neomycin, 500 µg/ml, Sigma Aldrich). Once the cells reached full confluence again, they were passaged, and again supplemented with the antibiotics. This cycle was repeated for at least three times a week for two weeks before initial use for production of rabies viral vectors. All cell lines presented in this study are available from the corresponding author upon request and all plasmids are available from Addgene (Supplementary Table 2).

### Generation of bicistronic CVS-N2c rabies viral vectors

A method to produce bicistronic reading frames in rabies viral vectors has previously been described for vectors of the SAD-B19 strain^11^. Here we adapted this approach for CVS-N2c vectors by inserting the virus’s endogenous N-P linker between two coding sequences, which led to efficient separation of the proteins without affecting the expression levels of either. The sequence for the linker used is: CATGAAAAAAActAACACTCCTCC (lower case letters indicate the N-P boundary).

### Production of rabies viral vectors

HEK293-GT cells were used to rescue both SAD-B19 and CVS-N2c rabies viral vectors. First, the cells were plated in a 35 mm culture dish and allowed to grow till they reached 80–90% confluence. Subsequently, they were transfected with the rabies vector plasmid and the SADB19 helper plasmids pTIT-N, pTIT-P and pTIT-L using polyethylenimine (PEI). Twenty-four hours later, the transfected cells were resuspended and re-plated in a 100 mm culture dish and incubated at 37 □C/5% CO_2_ until they regained full confluence. Cells were maintained that way with frequent medium changes until ∼100% of the cells were fluorescent, usually 5–6 days from time of transfection and 1–2 days from the point fluorescence was first detected. At this point the medium was harvested, filtered, aliquoted, and kept at −80 □C until further use.

For pseudotyping of rabies vectors, BHK-eT cells were used to simultaneously pseudotype and amplify the vectors: First, low confluence BHK-eT cells were plated in two 100 mm culture dishes and each transduced with 0.5 ml of the native-coat virus. Once the cells reached full confluence they were washed twice with DMEM, resuspended and each re-plated in a new 150mm culture dish. Once the cells reached full confluence again and ∼100% of them were fluorescent (∼3 days post transduction), the medium was collected, filtered, stored at 4 □C and replaced with fresh medium. This process was then repeated for two to three consecutive days. Following the last collection, the virus was pooled and centrifuged at 70,000 rcf for 1.5 hours. Following centrifugation, the medium was aspirated and the viral pellet resuspended in 200 µl phosphate-buffered saline (PBS), pH 7.4, aliquoted and stored at −80 □C until use.

For titration of envA-pseudotyped rabies viral vectors, HEK293-TVA were plated in 35 mm wells at low confluence along with one well containing HEK293T cells for detection of unpseudotyped particles. The following day, cells from one of the HEK293-TVA wells were resuspended and counted, while the cells in the remaining wells were transduced with 1 µl concentrated virus in serial dilutions ranging from 1:10–1:10000 while the HEK293T cells were transduced with 1 µl of undiluted virus. Three days later the cells were resuspended, washed with PBS and fixed with 4% PFA. The fraction of transduced cells in each well was determined using flow cytometry (FACS Aria III) and the titer was calculated based on a previously published formula^43^. In all of the titrations performed for viruses produced using our packaging system fluorescently-labeled cell were rarely detected in the control plate containing HEK293T cells, even when trasduced with vectors at concentrations an order of magnitude higher than required for complete labeling of HEK-TVA cells. All RVdG-CVS-N2c plasmids presented in this study are available from Addgene (Supplementary table 2).

### Production of adeno-associated viral vectors

Adeno-associated virus (AAV) production was performed in HEK293T cells based on a previously-published protocol^44^. Briefly, fully confluent HEK293 cells were transfected with an AAV2 vector plasmid along with pAdenoHelper and the AAV-dj RepCap plasmids using PEI. Thirty-six hours post transfection, the cells were harvested, pelleted, and lysed using three freeze-thaw cycles. The lysed cells were incubated with benzonase-nuclease (Sigma Aldrich) for one hour and then the debris was pelleted and the virus-containing supernatant collected and passed through a 0.22 µm filter. The collected supernatant was subsequently mixed with an equal amount of Heparin-agarose (Sigma Aldrich) and kept at 4 □C overnight with constant agitation. The following day, the agarose-virus mixture was transferred to a chromatography column and the agarose was allowed to settle. The supernatant was then drained from the column by means of gravity and the agarose-bound virus was washed once with PBS and then eluted using PBS supplemented with 0.5 M NaCl. The eluted virus was then filtered again, desalinated and concentrated using a 100 kDa centrifugal filter and then aliquoted and stored at −80 □C until use. All AAV plasmids presented in this study are available from Addgene (Supplementary Table 2).

### Animals

All transgenic driver and reporter lines have been previously characterized (Supplementary Table 3). In all experiments, male and female mice were used interchangeably in equal proportions, in an age range which varied between 1 to 6 months old. Neither sex nor age-related differences could be observed in any of the measurements. Experiments on C57BL/6 wild-type and transgenic mice were performed in strict accordance with institutional, national, and European guidelines for animal experimentation and were approved by the Bundesministerium für Wissenschaft, Forschung und Wirtschaft and Bildung, Wissenschaft und Forschung, respectively, of Austria (A. Haslinger, Vienna; BMWF-66.018/0010-WF/V/3b/2015; BMBWF-66.018/0008-WF/V/3b/2018).

### Stereotaxic intracranial virus injections

For *in vivo* delivery of viral vectors, 1 – 6 months old male or female mice were anaesthetized with isoflurane, injected with analgesics and placed in a stereotaxic frame where they continued to receive 1 – 5% isoflurane vaporized in oxygen at a fixed flow rate of 1 l min^-1^. Leg withdrawal reflexes were tested to evaluate the depth of anesthesia and when no reflex was observed an incision was made across the scalp to expose the skull. Bregma was then located and its coordinates used as reference for anterior-posterior (AP) and medio-lateral (ML) coordinates while the surface of the dura at the injection site was used as reference for dorso-ventral (DV) coordinates. In our experiments, we used the following sets of AP/ML/DV coordinates (in mm): DG: -1.9/1.3/-1.9; CA1: -1.9/1.5/-1.2; V1: -3.5/2/-0.7; LD: -1.3/1.3/-2.5. AAV vectors were first diluted 1:5 in PBS and delivered to the injection site at a volume of 0.3 µl and a rate of 0.06 µl per minute, using a Hamilton syringe and a 32G needle. After the injection was completed, the needle was left in place for a few additional minutes to allow the virus to diffuse in the tissue and then slowly retracted. At the end of the injection session, the scalp was glued back together and the mice were transferred back to their home cage to recover. Injection of pseudotyped rabies viral vectors took place 2–3 weeks after initial injection of AAV vectors containing the TVA receptor and rabies glycoprotein. Pseudotyped rabies vectors were first diluted to reach a final concentration of ∼2–5×10^8^ TU ml^-1^ and then injected in the same manner as the AAV. Except from rabies injections into the DG of Prox1cre transgenic animals, all other injections of rabies virus were shifted −0.2 mm AP and −0.2 mm ML. This was done in order to avoid, as much as possible, non-specific labeling along the needle tract of the first injection, due to the lack of complete specificity of cre-recombinase expression in the other transgenic lines used in this study.

### Slice preparation and electrophysiology

Electrophysiological recordings from identified retrogradely-labeled cells were performed 5–7 days following injection of CVS-N2c vectors. Manipulated animals were anaesthetized using an MMF mixture consisting of medetomidin (0.5 mg kg^-1^), midazolam (5 mg kg^-1^) and fentanyl (0.05 mg kg^-1^) and subsequently perfused through the heart with 20 ml ice cold dissection solution containing 87 mM NaCl, 25 mM NaHCO_3_, 2.5 mM KCl, 1.25 mM NaH_2_PO_4_, 10 mM D-glucose, 75 mM sucrose, 0.5 mM CaCl_2_, and 7 mM MgCl_2_ (pH 7.4 in 95% O_2_/5% CO_2_, 325 mOsm). The brain was then removed and the hippocampus along with the adjacent cortical tissue was dissected out and placed into a precast mold made of 4% agarose designed to stabilize the tissue. The mold was transferred to the chamber of a custom-built or a VT1200 vibratome (Leica Microsystems) and the tissue was transversely sectioned into 300 µm thick slices in the presence of ice-cold dissection solution. Transverse cortico-hippocampal sections were allowed to recover for ∼ 30 min at ∼ 31 °C and then kept at room temperature (20±1 °C) for the duration of the experiments. During recordings, slices were superfused with recording solution containing 125 mM NaCl, 2.5 mM KCl, 25 mM NaHCO_3_, 1.25 mM NaH_2_PO_4_, 25 mM D-glucose, 2 mM CaCl_2_, and 1 mM MgCl_2_ (pH 7.4 in 95% O_2_/5% CO_2_, ∼325 mOsm, at a rate of ∼1 ml min^-1^ using gravity flow. Labeled neurons in the regions of interest were identified according to their fluorescent signal and patched using pulled patch pipettes containing: 125 mM K-gluconate, 20 mM KCl, 0.1 mM EGTA, 10 mM phosphocreatine, 2 mM MgCl_2_, 2 mM ATP, 0.4 mM GTP, 10 mM HEPES (pH adjusted to 7.28 with KOH, ∼ 310 mOsm); 0.3% biocytin was added in a subset of recordings. Patched cells were maintained in current-clamp mode at the neuron’s resting membrane potential. Signals from patched cells were acquired using an Axon Axopatch 200A amplifier (Molecular Devices) and digitized using a CED Power 1401 analog-to-digital converter (Cambridge Electronic Design). Optogenetic stimulation was delivered using a blue-filtered white LED (Prizmatix, IL) at maximum intensity, passed through a 63x objective which was positioned above the recorded neuron. Ten bursts, spaced 20 seconds apart, consisting each of 5 light pulses (5 ms in duration) at 25 Hz were delivered to the tissue. Cells which exhibited responses with latency shorter than 1 ms or longer than 10 ms were excluded from the final analysis as they likely result from direct ChIEF activation in the recorded neuron or a polysynaptic response, respectively. Following these protocols, current steps ranging from −150 pA to 300 pA, with 25 pA increments, were delivered in current-clamp mode in order to determine firing frequency and input resistance, as previously described^45^. At the end of each recording session, the electrode was slowly retracted to create an outside-out patch, following which the slice was removed from the recording chamber, submerged in 4% PFA and then kept in 0.1 M phosphate buffer (PB) until further processing.

### Fixed tissue preparation

For imaging purposes, animals were sacrificed 5–7 days after injection of CVS-N2c vectors. First, animals were anaesthetized as described in the previous section and perfused through the heart with 15 ml of 0.1 PB followed by 30 ml of 4% PFA. Following perfusion, the brain was removed and kept in 4% PFA overnight at 4 □C, which was subsequently replaced with 0.1 M PB. Fixed brains were sectioned 100 µm thick in either a coronal, parasagittal or transverse plain and stored in 0.1 M PB at 4 □C.

For immunohistochemical labeling of transduced tissue, standard protocols were used. First, the sections were washed with PB 3X/10 min. Next, sections were incubated with 10% normal goat serum (NGS) and 0.4% Triton X-100 for 1 h, at room temperature (RT) with constant agitation and subsequently with the primary antibody (Rabbit anti somatostatin, 1:500, BMA Biomedicals; Rabbit anti parvalbumin, 1:1000, Swant antibodies; Chicken anti GFP, 1:000, Abcam; Mouse anti-FLAG, 1:1000, Sigma Alderich), in PB containing 5% NGS and 0.4% Triton X-100, at 4 □C overnight. After washing, slices were incubated with isotype-specific secondary antibodies (Alexa Fluor 647 conjugated goat anti rabbit, Alexa Fluor 647 conjugated goat anti mouse or Alexa Fluor 488 conjugated Goat anti-Chicken) in PB containing 5% NGS and 0.4% Triton X-100 for two hours in RT with constant agitation. After washing, slices were mounted, embedded in Prolong Gold Antifade mountant (Thermo-Fisher Scientific, Cat# P36930) and sealed with a 0.5 mm coverslip.

Neurons in acute slices that were filled with biocytin (0.3%) were processed for morphological analysis. After withdrawal of the pipettes, resulting in the formation of outside-out patches at the pipette tips, slices were fixed for 12–24 h at 4 °C in a 0.1 M PB solution containing 4% paraformaldehyde. After fixation, slices were washed, treated with Streptavidin Alexa Fluor 647 Conjugate (Thermo Fisher) for two hours at room temperature, washed again and embedded in Mowiol (Sigma-Aldrich).

### Microscopy and image analysis

All representative confocal images were acquired using either an LSM 800 microscope (Zeiss) or a Dragonfly spinning disc confocal microscope (Andor). All representative confocal images displayed in this manuscript were processed using FIJI^46^ and are shown as a maximal intensity projection of an image stack of 4-12 separate images. For quantification of cell numbers including channel overlap, Imaris 9 software was used.

### Cranial window implantation for *in vivo* CVS-N2c-GCaMP8m imaging

Before the surgery, animals were injected with Metacam (20 mg/kg s.c., 3.125 mg/ml solution) and Dexamethasone (0.2 mg kg^-1^ i.p., 0.02 mg ml^-1^ solution). Anesthesia was induced by 2.5% Isoflurane in oxygen in an anesthesia chamber. The mouse was subsequently fixed in a stereotaxic device (Kopf) with constant Isoflurane supply at 0.7 to 1.2% in O_2_ and body temperature controlled by a heating pad to 37.5 °C. After assertion that reflexes subsided, the cranium was exposed and cleaned of periost and connective tissue. A circular craniotomy of 4 mm diameter was drilled above V1, careful to leave the dura mater intact and the exposed brain constantly irrigated with artificial cerebrospinal fluid. A pulled glass capillary (tip diameter 30 to 40 µm) was loaded with CVS-N2c-GCaMP8m (5×10^8^ TU/ml) solution and 300 nl injected into the center of the craniotomy at a depth of 600 µm with a nanoliter injector (Nanoject, World Precision Instruments) at a speed of 30 nl min^-1^ and leaving the needle in place for 5 minutes after the volume was injected. Then, a 4 mm circular glass coverslip (CS-4R, Warner Instruments) was positioned on the brain and careful pressure applied with a toothpick mounted in the stereotaxic arm. The glass was first fixed in place with VetBond (3M). Then after cleaning and drying the surrounding cranium, a multilayer of glues was applied. First, to provide adhesion to the bone, All-in-One Optibond (Kerr) was applied and hardened by blue light (B.A. Optima 10). Second, Charisma Flow (Kulzer) was applied to cover the exposed bone and fix the glass in place by also applying blue light. After removal of the fixation toothpick, a custom designed and manufactured (RPD, Vienna) headplate, selective laser-sintered from the medical alloy TiAl6V4 (containing a small bath chamber and micro-ridges for repeatable fixation in the setup), was positioned in place and glued to the Charisma on the cranium with Paladur (Kulzer). Mice were given 300 µL of saline and 20 mg kg^-1^ metacam s.c., before removing them from the stereotaxic frame and letting them wake up while kept warm on a heating pad. Another dose of 20 mg/kg metacam s.c. and 0.2 mg kg^-1^ i.p. Dexamethasone was further injected 24 hours after conclusion of the surgery.

### Setup and visual stimuli

Mice were head-fixed using a custom-manufactured clamp that was connected to a 3-axis motorized stage (8MT167-25LS, Standa). Mice could run freely on a custom-designed spherical treadmill (20 cm diameter). Visual stimuli were projected by a modified LightCrafter (Texas Instruments) at 60 Hz, reflected by a quarter-sphere mirror (Modulor) below the mouse and presented on a custom-made spherical dome (80 cm diameter) with the mouse’s head at its center. The green and blue LEDs in the projector were replaced by cyan (LZ1-00DB00-0100, Osram) and UV (LZ1-00UB00-01U6, Osram) LEDs respectively. A double band-pass filter (387/480 HD Dualband Filter, Semrock) was positioned in front of the projector to not contaminate the imaging. The reflected red channel of the projector was captured by a transimpedance photo-amplifier (PDA36A2, Thorlabs) and digitized for synchronization. Cyan and UV LED powers were adjusted to match the relative excitation of M-and S-cones during an overcast day, determined and calibrated using opsin templates^47^ and a spectrometer (CCS-100, Thorlabs). Stimuli were designed and presented with Psychtoolbox^48^, running on MATLAB 2020b (Mathworks). Stimulus frames were morphed on the GPU using a customized projection map and an OpenGL shader to counteract the distortions resulting from the spherical mirror and dome. The dome setup allows to present mesopic stimuli from ca. 90° on the left to ca. 170° on the right in azimuth and from ca. 40° below to ca. 80° above the equator in elevation. During anatomical stack imaging, dense moving dots of different sizes and light intensities, moving in uncorrelated directions were shown, to excite neurons with a complex texture-like stimulus. For functional imaging, full field step gratings with temporal frequency of 2 Hz and spatial frequency of 0.1 cycles/° were shown moving in 8 randomly ordered directions. In each trial, the grating image remained stationary for 3 seconds and then moved for 7 seconds in the respective direction. Each direction was shown 5-10 times in total per session.

### Imaging

Two-photon imaging was performed on a custom-built microscope, controlled by Scanimage (Vidrio Technologies) running on MATLAB 2020b (Mathworks) and a PXI system (National Instruments). The beam from a pulsed Ti:Sapphire laser (Mai-Tai DeepSee, Spectra-Physics) was scanned by a galvanometric-resonant (8 kHz) mirror combination (Cambridge Scientific) and expanded to underfill the back-aperture of the objective (16x 0.8 N.A. water-immersion, Nikon); 1.9 by 1.9 mm field-of-view; 30 Hz frame rates. Fast volumetric imaging was acquired with a piezo actuator (P-725.4CA, Physik Instrumente). Emitted light was collected (FF775-Di01, Semrock), split (580 nm long-pass, FF580-FDi01, Semrock), band-pass filtered (green: FF03-525/50; red: FF01-641/75, Semrock), measured (GaAsP photomultiplier tubes, H10770B-40, Hamamatsu), amplified (TIA60, Thorlabs) and digitized (PXIe-7961R NI FlexRIO FPGA, NI 5734 16-bit, National Instruments). Laser wavelength was set to 935 or 955 nm, which excited GCaMP8 well and tdTomato sufficiently for anatomical identification. Maximum laser power used at the deepest planes was 80 mW/mm^2^. Due to the early start of imaging after implantation the tissue cleared only over the course of the imaging days, necessitating relatively high laser powers in the beginning. To avoid heat damage, only 15 minutes of continuous imaging was performed, after which imaging was paused for at least 5 minutes^49^. At each recording day (day 3, 5, 7, 9, 11, 14, 16 post RVdG_envA-_CVS-N2c-GCaMP8m injection), first a dense anatomical stack with a 10 µm plane distance over the full accessible depth and plane averaging over 25 frames was recorded in the injected area. If GCaMP labeled neurons were found, subsequently a functional imaging session was started over 5-8 z-planes with 25-50 µm plane distance and voxel size of 1.4 to 1.7 µm resulting in a volume rate of 4.2 to 5 Hz.

### Imaging data analysis

Cell numbers were estimated by Imaris (Oxford Instruments) on anatomical stack images. Functional calcium imaging data was first analyzed with suite2p (v0.10.0)^50^ for motion correction and ROI extraction. ROIs were then curated manually based on morphological and activity shape. Further analysis was performed in custom MATLAB R2021a (Mathworks) scripts: dF/F0 was estimated based on published procedures^51^ by first subtracting neuropil contamination (from suite2p, fluorescence signal of 350 pixels surrounding the ROI, excluding other ROIs) with a factor of 0.5 (estimated from fluorescence of small capillaries as reported previously ^52^. From the neuropil-corrected ROI fluorescence, baseline F0 was defined as the 8th percentile of a moving window of 15 seconds ^53^. dF/F0 was then calculated on the same window by first subtracting and then dividing fluorescence trace by median of the same 15 second window^51^. Signal-to-Noise ratio (SNR) was defined for each neuron by dividing the 99th percentile of the dF/F trace (“signal”) by the standard deviation of its negative values after baseline correction (“noise”). Direction selectivity index (DSI) and preferred direction was calculated based on the vector sum method ^54^on the mean dF/F0 of the 7 seconds per direction the grating was moving. DSI significance was estimated by a permutation test of the direction labels (resampled 1000 times) to define the proportion of DSIshuffled>DSI.

### Statistical analysis

All values were reported as mean and error bars as ± SEM. Statistical significance was tested using non-parametric, double-sided Kruskal-Wallis test followed by a double-sided Mann-Whitney test for post-hoc comparisons, or by Fisher’s exact test, in Microsoft Excel. Differences with p < 0.05 were considered significant. In figures, a single asterisk (*), double asterisks (**), and triple asterisks (***) indicate p < 0.05, p < 0.01 and p < 0.001, respectively.

## Acknowledgements

We would like to thank F. Marr for technical assistance, to A. Murray for RVdG-CVS-N2c viruses and Neuro2A packaging cell-lines and to J. Watson for reading the manuscript. This research was supported by the Scientific Service Units (SSU) of IST-Austria through resources provided by the Bioimaging Facility (BIF) and the Preclinical Facility (PCF). This project was funded by the European Research Council (ERC) under the European Union’s Horizon 2020 research and innovation programme (ERC advanced grant No 692692) and the Fond zur Förderung der Wissenschaftlichen Forschung (Z 312-B27, Wittgenstein award).

## Author contributions

Y.B. designed and performed all experiments, A.S., M. J. and Y.B. designed calcium imaging experiments, which were performed and analyzed by A.S. P.J. supervised the project, Y.B. and A.S. wrote the manuscript, with comments from all authors.

---

**Extended Data Figure. 1.**
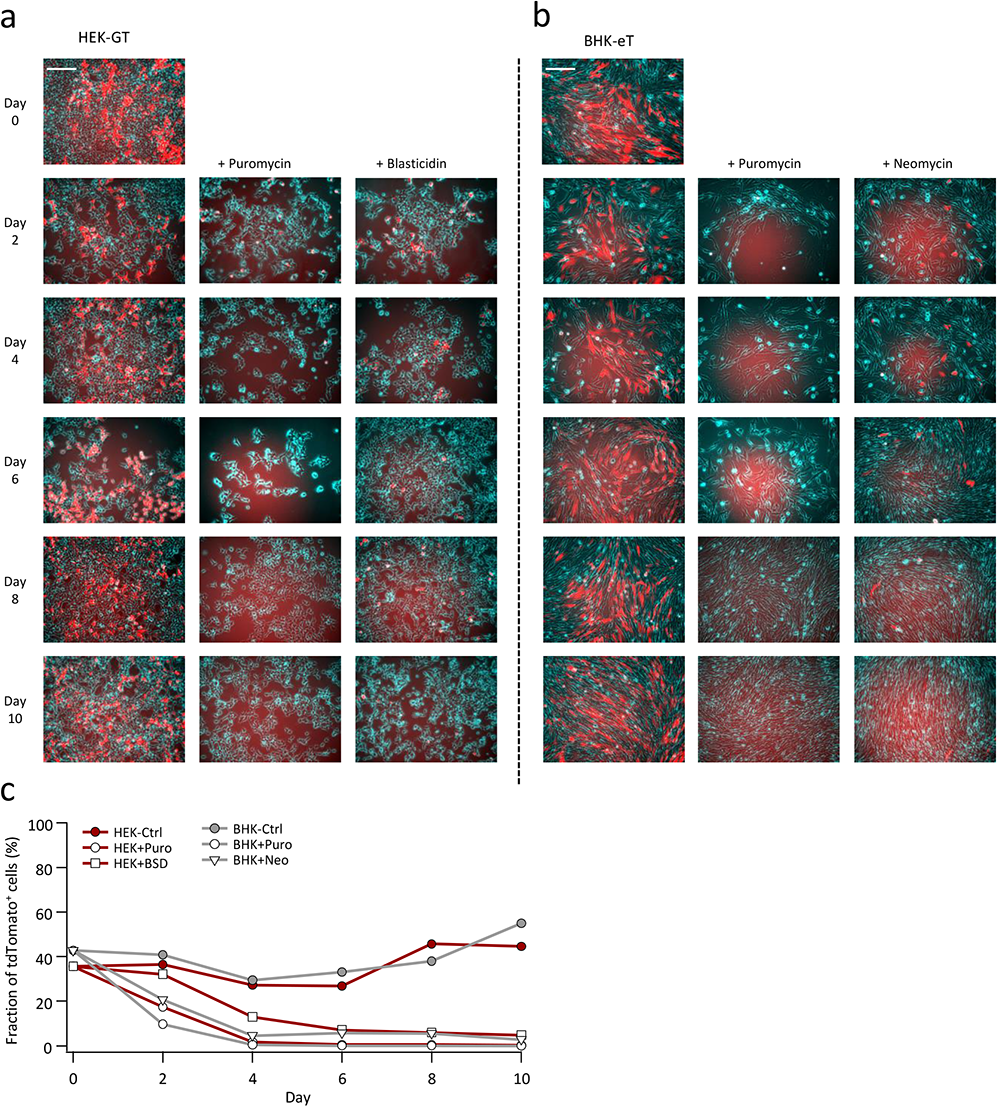
Efficiency of stable cell selection using antibiotic resistance genes. **a,b,** HEK-GT (A) and BHK-eT (B) cells (bright-field illumination shown in cyan) were mixed with HEK293 or BHK-21 cells, respectively, stably expressing tdTomato only (red). Representative images show the gradual removal of tdTomato+ cells following exposure to either of the antibiotics Puromycin, Blasticidin or used in the generation of the stable cells. Scale bars represent 100 µm. **c,** FACS-assisted quantification of the fraction of tdTomato^+^ cells shows the time course for their removal from the antibiotics-exposed cultures.

**Extended Data Figure 2.**
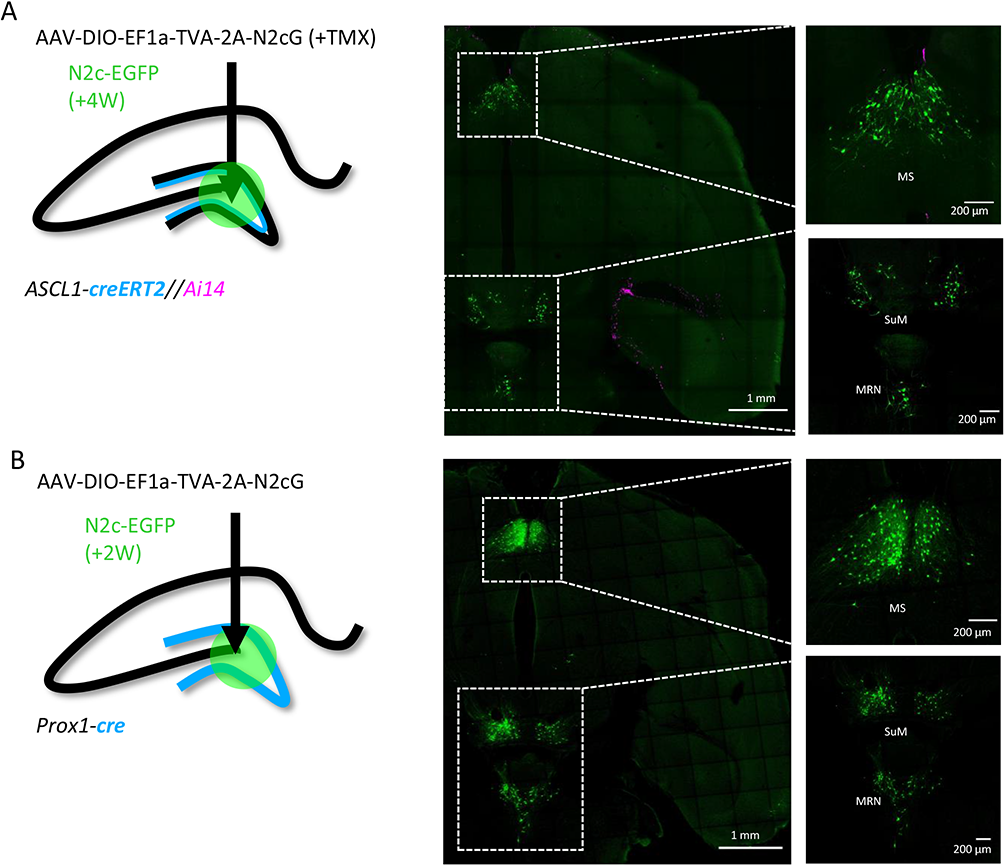
Trans synaptic retrograde labeling pattern from newborn hippocampal DGCs in subcortical regions. **a,b,** Transsynaptic retrograde labeling with CV-N2c vectors from adult-born cells in the hippocampal DG, using the Ascl1-creERT2 line (a), reveals identical labeling pattern in subcortical regions as following the same manipulation on the general population of DGCs, using the DG-specific Prox1cre line (b). MS – Medial Septal Nucleus; SuM – Supramammilary Nucleus; MRN – Median Raphe nucleus.

**Extended Data Figure 3.**
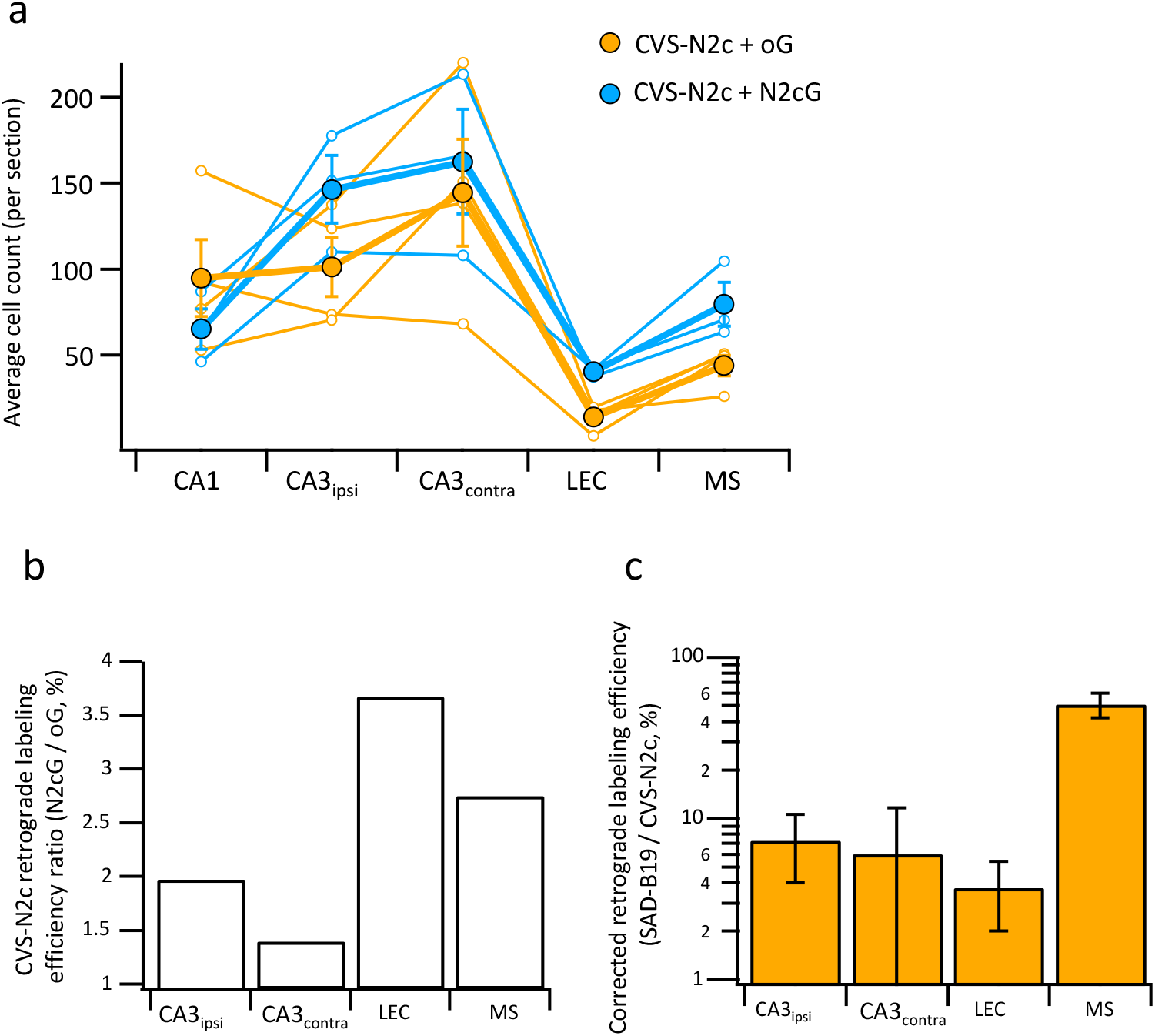
Differential effects of N2c and B19 glycoproteins on propagation efficiency of CVS-N2c vectors. **a,** Average neuron count per slice for individual animals (empty circles) and averaged count for all animals (full circles) in the different regions projecting to the CA1 following retrograde labeling with CVS-N2c vectors using either the B19-oG (Orange) or the N2cG (Cyan). **b,** Ratio of 2^nd^ order neurons between the two conditions in (a), normalized by the number of putative starter neurons in the CA1 pyramidal layer demonstrates higher efficacy of propagation under the N2cG, for all regions tested. **c,** Retrograde labeling efficacy ratio between CVS-N2c and SAD-B19 vectors shown in Figure 2h, corrected for the added effect of the glycoprotein to the propagation efficacy of the CVS-N2c vectors by dividing the values for the SAD-B19 condition in Figure 2h, with the corresponding ratio estimates measured in (b).

**Extended Data Figure 4.**
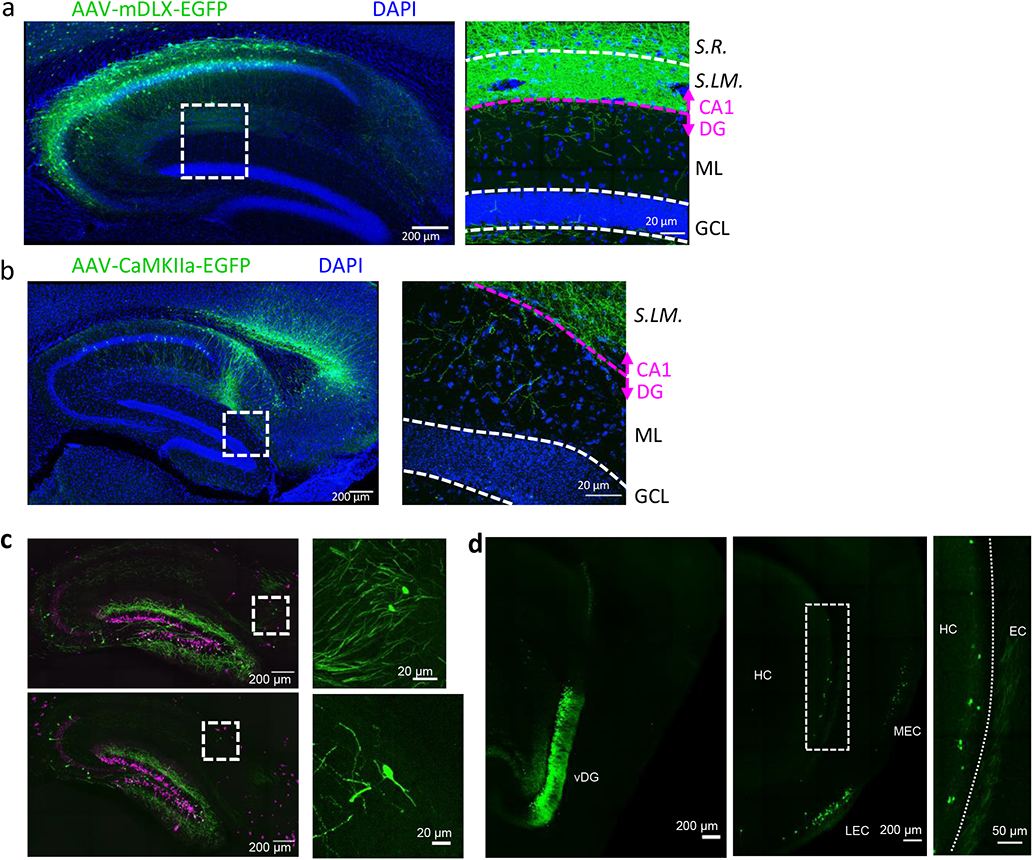
Cross validation of novel excitatory and inhibitory intrahippocampal projections to DGCs. **a,b,** Both injection of AAV-mDLX-EGFP, for specific expression in inhibitory neurons, into the CA1 (a), as well as injection of AAV-CaMKII-EGFP, for specific expression in excitatory neurons, into the subiculum (b) reveal axonal arborizations in the molecular layer of the DG. **c,** Retrograde labeling from a sparse population of adult-born DGCs (as shown in Figure 1j) also reveals projections from the subiculum (top) and S.O. (bottom). **d,** Retrograde labeling from the ventral DG (vDG) using the Prox1-cre line (left panel) shows projection neurons along the superficial most layer of the ventral subiculum (middle and right panels)

**Extended Data Figure 5.**
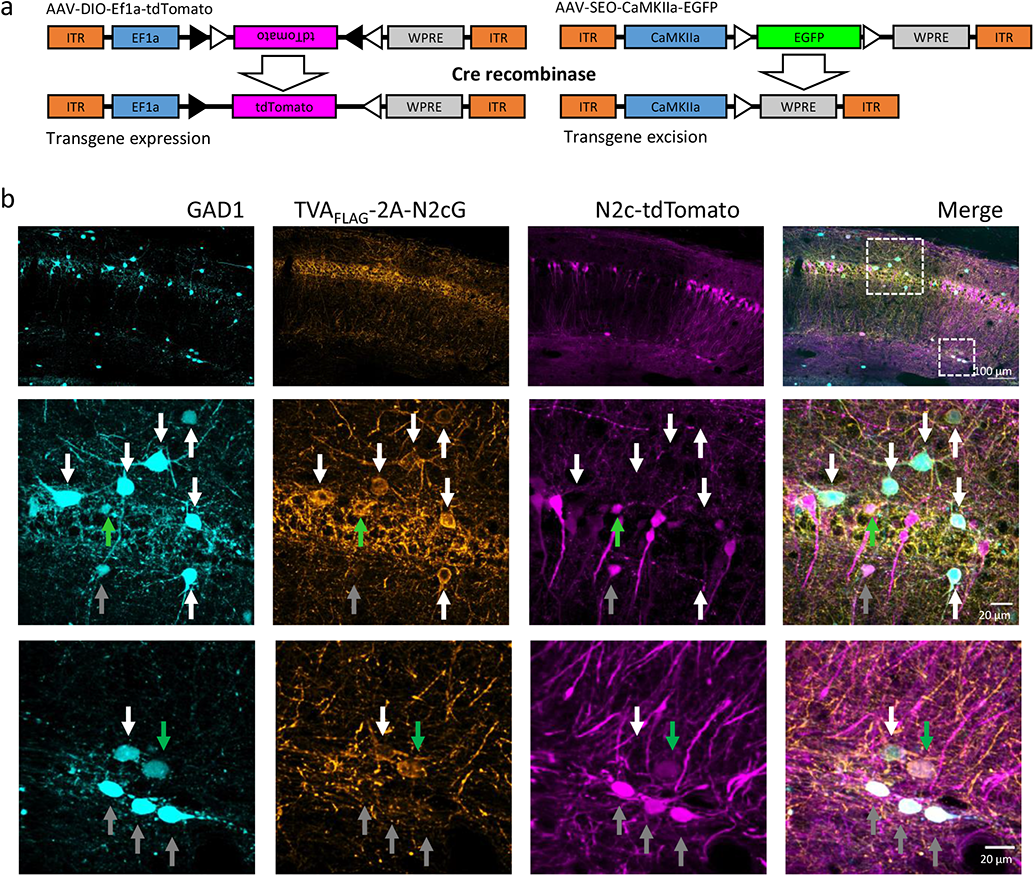
Targeting specificity of newly-designed AAV vectors. **a,** A schematic illustration demonstrating the differential effect of cre-mediated recombination on double-floxed, inverted open reading frames (DIO, left) or single-floxed, excisable open reading frames (SEO, right). **b,** Representative confocal images from the hippocampus of a GAD1-EGFP mouse (Cyan), injected with AAV-mDLX-TVA-2A-N2cG (orange) and subsequently CVS-N2c-tdTomato (magenta) into the hippocampal CA1. Green arrows in the expanded images indicate starter neurons expressing all three markers, grey arrows indicate 2^nd^ order interneurons, expressing EGFP and tdTomato, but not TVA_FLAG_-2A-N2cG, and white arrows indicate potential, yet non-participating interneuorons expressing GAD1 and TVA_FLAG_-2A-N2cG, but not tdTomato.

**Extended Data Figure 6.**
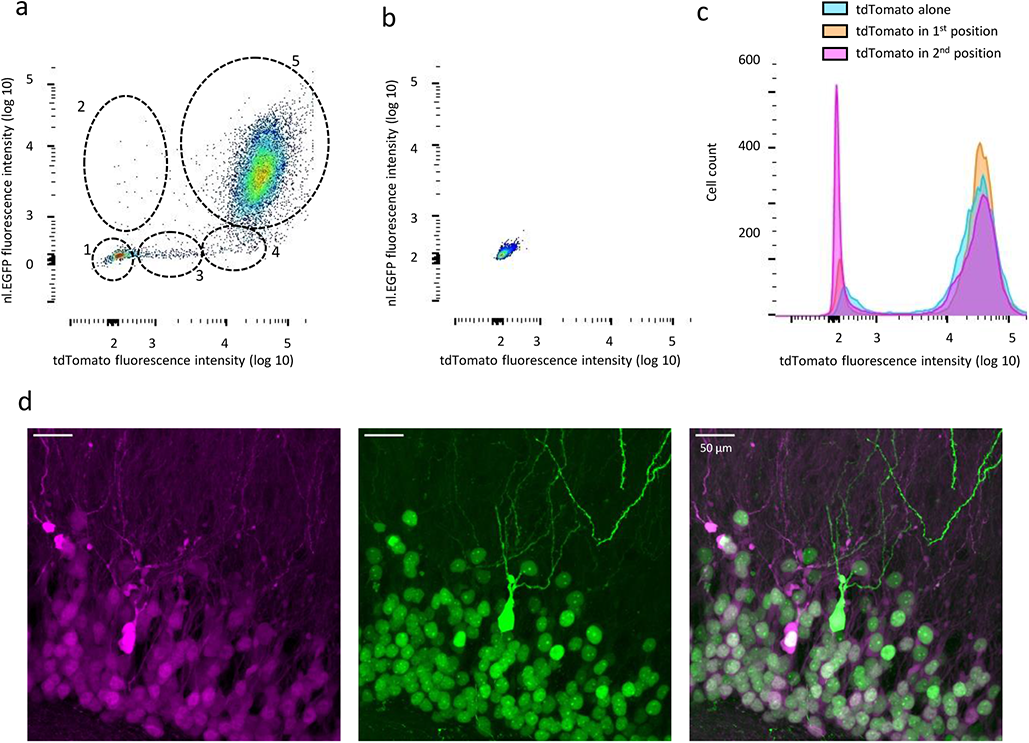
Additional properties of bicistronic RVdG-CVS-N2c vectors. **a,b,** FACS analysis of fluorescence intensity in HEK-TVA cells transduced with RVdG_envA_-CVS-N2c-nl.EGFP-tdTomato (a) or HEK293 cells transduced with 10X concentration of the same vector (b). Five different clusters of labeled cells are highlighted in (a), likely representing: 1) tdTomato^-^/nl.EGFP^-^ Non-transduced cells; 2) tdTomato^-^/nl.EGFP^+^ Cells with a putative null mutation in the tdTomato gene; 3) tdTomato^low^/nl.EGFP^-^ putatively undergoing mytosis and lack a defined nucleus,; 4) tdTomato^+^/nl.EGFP^-^ with a putative null mutation in the EGFP gene; 5) tdTomato^+^/nl.EGFP^+^ with no putative mutations. **c,** Quantification of fluorescence intensity for three different RVdG-CVS-N2c vectors, in which tdTomato is expressed alone, or in a bicistronic vector in either the first (tdTomato-FlpO) or second (nl.EGFP-tdTomato) position shows no substantial differences in tdTomato fluorescence intensity. **d,** A representative, high-magnification confocal image of the granular cell layer of the DG, following targeting of the RVdG_envA_-CVS-N2c-nl.EGFP-tdTomato vector, showing a cell expressing a soluble EGFP, which is likely the result of a mutation in the nuclear localization signal.

**Extended Data Figure 7.**
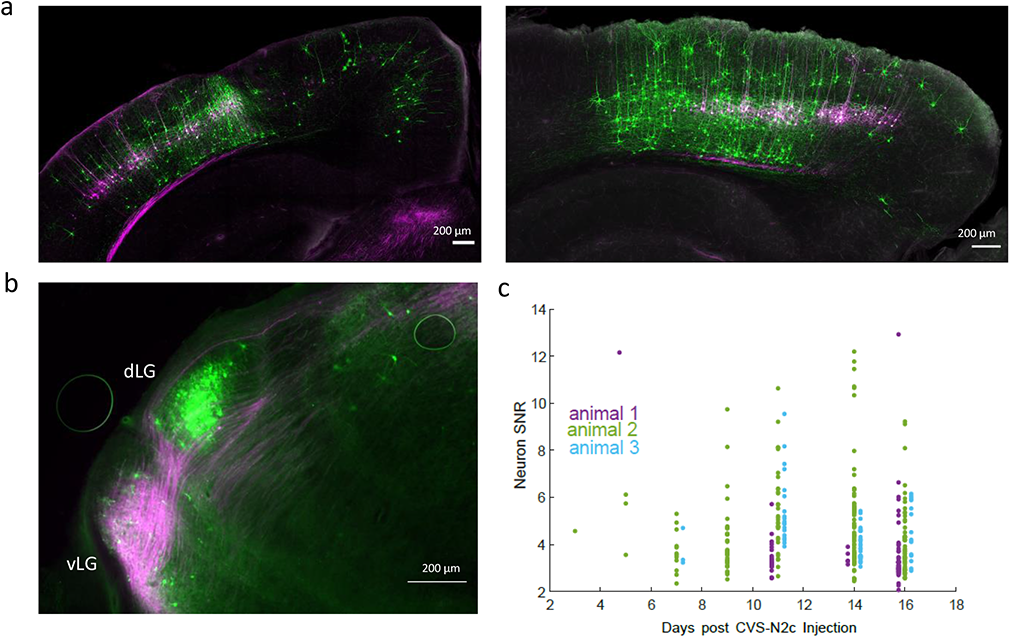
Retrograde labeling specificity for in-vivo calcium imaging with RVdG_envA_-CVS-N2c-GCaMP8m. **a,** Representative confocal images of the V1 area in the coronal plane (animal 2, left) or of the sagittal plane (animal 3, right) of the two additional animals used in the experiment. **b,** A representative fluorescent microscope image of the lateral geniculate nucleus showing retrogradely-labeled, V1 layer 5-projecting neurons in the dorsolateral geniculate nucleus (dLG) and fibers originating in V1 layer V neurons in the ventrolateral geniculate nucleus (vLG). **c,** A time plot showing the gradual change in signal-to-noise ratio (SNR) for all three animals throughout the experiment.

## References

1. Fornito, A., Zalesky, A. & Breakspear, M. The connectomics of brain disorders. Nature Reviews Neuroscience vol. 16 159–172 (2015).

2. Morgan, J. L. & Lichtman, J. W. Why not connectomics? Nature Methods vol. 10 494–500 (2013).

3. Luo, L., Callaway, E. M. & Svoboda, K. Genetic dissection of neural circuits. Neuron 57, 634– 660 (2008).

4. Luo, L., Callaway, E. M. & Svoboda, K. Genetic Dissection of Neural Circuits: A Decade of Progress. Neuron vol. 98 256–281 (2018).

5. Tervo, D. G. R. et al. A designer AAV variant permits efficient retrograde access to projection neurons. Neuron 92, 372–382 (2016).

6. Zingg, B. et al. AAV-Mediated Anterograde Transsynaptic Tagging: Mapping Corticocollicular Input-Defined Neural Pathways for Defense Behaviors. Neuron 93, 33–47 (2017).

7. Zampieri, N., Jessell, T. M. & Murray, A. J. Mapping sensory circuits by anterograde transsynaptic transfer of recombinant rabies virus. Neuron 81, 766–778 (2014).

8. Ginger, M., Haberl, M., Conzelmann, K. K., Schwarz, M. K. & Frick, A. Revealing the secrets of neuronal circuits with recombinant rabies virus technology. Frontiers in Neural Circuits vol. 7 2 (2013).

9. Wickersham, I. R., Finke, S., Conzelmann, K. K. & Callaway, E. M. Retrograde neuronal tracing with a deletion-mutant rabies virus. Nature Methods 4, 47–49 (2007).

10. Albisetti, G. W. et al. Identification of two classes of somatosensory neurons that display resistance to retrograde infection by rabies virus. Journal of Neuroscience 37, 10358–10371 (2017).

11. Osakada, F. & Callaway, E. M. Design and generation of recombinant rabies virus vectors. Nat Protoc 8, 1583–601 (2013).

12. Reardon, T. R. et al. Rabies virus CVS-N2cδG strain enhances retrograde synaptic transfer and neuronal viability. Neuron 89, 711–724 (2016).

13. Norrman, K. et al. Quantitative comparison of constitutive promoters in human ES cells. PLoS ONE 5, (2010).

14. Borges-Merjane, C., Kim, O. & Jonas, P. Functional Electron Microscopy, “Flash and Freeze,” of Identified Cortical Synapses in Acute Brain Slices. Neuron 105, 992–1006 (2020).

15. Yang, S. M., Alvarez, D. D. & Schinder, A. F. Reliable genetic labeling of adult-born dentate granule cells using ascl1CreERT2 and glastCreERT2 murine lines. Journal of Neuroscience 35, 15379–15390 (2015).

16. Deshpande, A. et al. Retrograde monosynaptic tracing reveals the temporal evolution of inputs onto new neurons in the adult dentate gyrus and olfactory bulb. Proc Natl Acad Sci U S A 110, 1152–1161 (2013).

17. Vivar, C. et al. Monosynaptic inputs to new neurons in the dentate gyrus. Nature Communications 3, 1107 (2012).

18. Rowland, D. C. et al. Transgenically targeted rabies virus demonstrates a major monosynaptic projection from hippocampal area CA2 to medial entorhinal layer II neurons. Journal of Neuroscience 33, 14889–14898 (2013).

19. Kim, E. J., Jacobs, M. W., Ito-Cole, T. & Callaway, E. M. Improved Monosynaptic Neural Circuit Tracing Using Engineered Rabies Virus Glycoproteins. Cell Reports 15, 692–699 (2016).

20. Hájos, N. & Mody, I. Synaptic communication among hippocampal interneurons: Properties of spontaneous IPSCs in morphologically identified cells. Journal of Neuroscience 17, 8427– 8442 (1997).

21. Katona, L. et al. Behavior-dependent activity patterns of GABAergic long-range projecting neurons in the rat hippocampus. Hippocampus 27, 359–377 (2017).

22. Klausberger, T. & Somogyi, P. Neuronal diversity and temporal dynamics: The unity of hippocampal circuit operations. Science (1979) 321, 53–57 (2008).

23. Szabo, G. G. et al. Extended Interneuronal Network of the Dentate Gyrus. Cell Reports 20, 1262–1268 (2017).

24. Tamamaki, N. et al. Green fluorescent protein expression and colocalization with calretinin, parvalbumin, and somatostatin in the GAD67-GFP knock-in mouse. Journal of Comparative Neurology 467, 60–79 (2003).

25. Freund, T. F. & Buzsáki, G. Interneurons of the Hippocampus. Hippocampus 6, 347–470 (1996).

26. Li, Y. et al. A distinct entorhinal cortex to hippocampal CA1 direct circuit for olfactory associative learning. Nature Neuroscience 20, 559–570 (2017).

27. Valero, M. et al. Determinants of different deep and superficial CA1 pyramidal cell dynamics during sharp-wave ripples. Nature Neuroscience 18, 1281–1290 (2015).

28. Masurkar, A. V. et al. Medial and Lateral Entorhinal Cortex Differentially Excite Deep versus Superficial CA1 Pyramidal Neurons. Cell Reports 18, 148–160 (2017).

29. Dimidschstein, J. et al. A viral strategy for targeting and manipulating interneurons across vertebrate species. Nature Neuroscience 19, 1743–1749 (2016).

30. Basu, J. et al. Gating of hippocampal activity, plasticity, and memory by entorhinal cortex long-range inhibition. Science (1979) 351, aaa5694 (2016).

31. Melzer, S. et al. Long-range-projecting gabaergic neurons modulate inhibition in hippocampus and entorhinal cortex. Science (1979) 335, 1506–1510 (2012).

32. Madisen, L. et al. Transgenic Mice for Intersectional Targeting of Neural Sensors and Effectors with High Specificity and Performance. Neuron 85, 942–958 (2015).

33. Gloveli, T., Dugladze, T., Schmitz, D. & Heinemann, U. Properties of entorhinal cortex deep layer neurons projecting to the rat dentate gyrus. European Journal of Neuroscience 13, 413– 420 (2001).

34. Osakada, F. et al. New rabies virus variants for monitoring and manipulating activity and gene expression in defined neural circuits. Neuron 71, 617–631 (2011).

35. Lin, J. Y., Lin, M. Z., Steinbach, P. & Tsien, R. Y. Characterization of engineered channelrhodopsin variants with improved properties and kinetics. Biophysical Journal 96, 1803–1814 (2009).

36. Chatterjee, S. et al. Nontoxic, double-deletion-mutant rabies viral vectors for retrograde targeting of projection neurons. Nature Neuroscience 21, 638–646 (2018).

37. Rossi, L. F., Harris, K. D. & Carandini, M. Spatial connectivity matches direction selectivity in visual cortex. Nature 588, 648–652 (2020).

38. Wertz, A. et al. Single-cell-initiated monosynaptic tracing reveals layer-specific cortical network modules. Science (1979) 349, 70–74 (2015).

39. Wickersham, I. R. et al. Monosynaptic restriction of transsynaptic tracing from single, genetically targeted neurons. Neuron 53, 639–647 (2007).

40. Ohara, S. et al. Intrinsic Projections of Layer Vb Neurons to Layers Va, III, and II in the Lateral and Medial Entorhinal Cortex of the Rat. Cell Reports 24, 107–116 (2018).

41. Chattopadhyaya, B. et al. Experience and activity-dependent maturation of perisomatic GABAergic innervation in primary visual cortex during a postnatal critical period. Journal of Neuroscience 24, 9598–9611 (2004).

42. Zhang, Y., et al. Fast and sensitive GCaMP calcium indicators for imaging neural populations. bioRxiv (2021).

43. Wickersham, I. R., Sullivan, H. A. & Seung, H. S. Production of glycoprotein-deleted rabies viruses for monosynaptic tracing and high-level gene expression in neurons. Nature Protocols 5, 595–606 (2010).

44. McClure, C., Cole, K. L. H., Wulff, P., Klugmann, M. & Murray, A. J. Production and titering of recombinant adeno-associated viral vectors. J Vis Exp 57, (2011).

45. Kowalski, J., Gan, J., Jonas, P. & Pernia-Andrade, A. J. Intrinsic membrane properties determine hippocampal differential firing pattern in vivo in anesthetized rats. Hippocampus 26, 668–682 (2016).

46. Schindelin, J., et al. Fiji: An open-source platform for biological-image analysis. Nature Methods vol. 9 676–682 (2012).

47. Govardovskii, V. I., Fyhrquist, N., Reuter, T., Kuzmin, D. G. & Donner, K. In search of the visual pigment template. Visual Neuroscience 17, 509–528 (2000).

48. Brainard, D. H. The Psychophysics Toolbox. Spatial Vision 10, 433–436 (1997).

49. Podgorski, K. & Ranganathan, G. Brain heating induced by near-infrared lasers during multiphoton microscopy. Journal of Neurophysiology 116, 1012–1023 (2016).

50. Pachitariu, M. et al. Suite2p: beyond 10,000 neurons with standard two-photon microscopy. bioRxiv 061507 (2016) doi:10.1101/061507.

51. Keller, G. B., Bonhoeffer, T. & Hübener, M. Sensorimotor Mismatch Signals in Primary Visual Cortex of the Behaving Mouse. Neuron 74, 809–815 (2012).

52. Kerlin, A. M., Andermann, M. L., Berezovskii, V. K. & Reid, R. C. Broadly Tuned Response Properties of Diverse Inhibitory Neuron Subtypes in Mouse Visual Cortex. Neuron 67, 858– 871 (2010).

53. Dombeck, D. A., Khabbaz, A. N., Collman, F., Adelman, T. L. & Tank, D. W. Imaging Large-Scale Neural Activity with Cellular Resolution in Awake, Mobile Mice. Neuron 56, 43–57 (2007).

54. Mazurek, M., Kager, M. & Van Hooser, S. D. Robust quantification of orientation selectivity and direction selectivity. Frontiers in Neural Circuits 8, 92 (2014).

