## Supplementary tables for "Fast, high-throughput production of improved rabies viral vectors for specific, efficient and versatile transsynaptic retrograde labeling"

Supplementary Table 1: Comparison of packaging systems

**Rescue from DNA and amplification of native-coat stock**

|  | B7GG^1^ | Neuro2a-N2cG^2^ | HEK-GT |
| --- | --- | --- | --- |
| Stably-expressed transgenes | T7 polymerase + SAD-B19G | CVS-N2cG | Optimized T7 polymerase (oT7) + optimized SAD-B19G (oG) |
| Selection markers | Fluorescence | Fluorescence | Antibiotic resistance genes |
| Transfected genes | Vector + N,P, G & L | Vector + T7, N,P,G & L | Vector + N,P & L |
| Transfection efficiency | Low | Low | High |
| Growth conditions | 3% CO2 at 35°C | 3% CO2 at 35°C | 5% CO2 at 37°C |
| Rescue timeline | 10-11 days | 10-11 days | 5-6 days |
| Initial amplification timeline | 9-11 days | 14-21 days | Not required |
| Compatibility | SAD-B19 (CVS-N2c possible, but not tested | CVS-N2c only | Both SAD-B19 and CVS N2c |

**Pseudotyping**

|  | BHK-EnvA^1^ | Neuro2a-envA^2^ | BHK-eT |
| --- | --- | --- | --- |
| Stably-expressed transgenes | envA or envB | envA | envA + TVA |
| Selection markers | Fluorescence | Fluorescence | Antibiotic resistance genes |
| Growth conditions | 3% CO2 at 35°C | 3% CO2 at 35°C | 5% CO2 at 37°C |
| Pseudotyping timeline | 7-10 days | 28 days | 4-6 days |
| Requirements for pseudotyping | Large stock of concentrated native-coat particles | Large stock of concentrated native-coat particles | Trace amounts of either native-coat or pseudotyped stock |
| Titer | Low 10^8 typical | Low 10^7 typical | High 10^9 typical |
| Native-coat background | 10^2 typical | Not detectable | Not detectable |

1.	Osakada, F. & Callaway, E. M. Design and generation of recombinant rabies virus vectors. *Nature protocols* **8**, 1583–601 (2013).

2.	Reardon, T. R. *et al.* Rabies virus CVS-N2cδG strain enhances retrograde synaptic transfer and neuronal viability. *Neuron* **89**, 711–724 (2016).

Supplementary Table 2: Summary of plasmids.

| **Plasmid name** | **Source (deposited by)** | **Cat#** |
| --- | --- | --- |
| pCAG-B19N | AddGene (I. Wickersham) | #59924 |
| pCAG-B19P | AddGene (I. Wickersham) | #59925 |
| pCAG-B19L | AddGene (I. Wickersham) | #59922 |
| pAdDeltaF6 | AddGene (J. Wilson) | #112867 |
| rAAV-DJ RepCap | Gift from Mark A. Kay |  |
| rAAV2-retro helper | AddGene (A. Karpova and D. Schaffer) | #81070 |
| pAAV-EF1a-Cre | AddGene (K. Deisseroth) | #55636 |
| pAAV-DIO-hSyn-mCherry | AddGene (K. Deisseroth) | #114472 |
| RVdG-CVS-N2c-EGFP | AddGene (T. Jessell) | #73461 |
| RVdG-CVS-N2c-tdTomato | AddGene (T. Jessell) | #73462 |
| pAAV-DIO-Ef1a-TVA-2A-oG | This paper | #172359 |
| pAAV-DIO-Ef1a-TVA-2A-N2cG | This paper | #172360 |
| pAAV-FRT-EF1a-TVA-2A-N2cG | This paper | #172361 |
| pAAV-DIO-CaMKII-TVA-P2A-N2cG | This paper | #172362 |
| pAAV-SEO-CaMKII-TVA-P2A-N2cG | This paper | #172363 |
| pAAV-mDLX-TVA-2A-N2cG | This paper | #172364 |
| pAAV-DIO-mDLX-TVA-2A-N2cG | This paper | #172365 |
| pAAV-DIO-CAG-TVA-P2A-dTomato | This paper | #177016 |
| pAAV-DIO-EF1a-TVA-P2A-EYFP | This paper | #177017 |
| pAAV-SEO-CaMKII-EGFP | This paper | #177018 |
| MMLV-CAG-TVA-IRES-Puro | This paper | #172366 |
| MMLV-CAG-SADB19_oG-IRES-Puro | This paper | #172367 |
| MMLV-CAG-G_oT7pol-IRES-BSD | This paper | #172369 |
| pLV-EF1a-N2c_envA-IRES-Neo | This paper | #172368 |
| CVS-N2c-tdTomato-ChIEF | This paper | #172370 |
| CVS-N2c-EGFP-ChIEF | This paper | #172371 |
| CVS-N2c-EGFP-iCre | This paper | #172372 |
| CVS-N2c-EGFP-FlpO | This paper | #172373 |
| CVS-N2c-tdTomato-iCre | This paper | #172374 |
| CVS-N2c-tdTomato-FlpO | This paper | #172375 |
| CVS-N2c-mTurquoise | This paper | #172376 |
| CVS-N2c-E2_Crimson | This paper | #172377 |
| CVS-N2c-nl.mCherry-FlpO | This paper | #172378 |
| CVS-N2c-nl.EGFP-FlpO | This paper | #172379 |
| CVS-N2c-nl.EGFP-SypGFP | This paper | #172380 |
| CVS-N2c-SypRFP | This paper | #172381 |
| CVS-N2c-nl.EGFP-tdTomato | This paper | #172382 |
| CVS-N2c-EYFP | This paper | #172383 |
| CVS-N2c-mCitrine | This paper | #172384 |
| CVS-N2c-nl.mCherry-GCaMP7s | This paper | #172385 |
| CVS-N2c-nl.EGFP-jRGECO1a | This paper | #172386 |
| CVS-N2c-GCaMP8f | This paper | #172387 |
| CVS-N2c-GCaMP8m | This paper | #172388 |
| CVS-N2c-GCaMP8s | This paper | #172389 |

Supplementary Table 3: Summary of transgenic lines.

| Transgenic line | Full name | Source (deposited by) | Cat# |
| --- | --- | --- | --- |
| Prox1^cre^ | Tg(Prox1-cre)SJ32Gsat/Mmucd | MMRRC (N. Heintz) | 036644-UCD |
| Calb1-cre | B6;129S-*Calb1^tm2.1(cre)Hze^*/J | Jackson labs (H. Zeng) | 028532 |
| Ascl1-cre | *Ascl1^tm1.1(Cre/ERT2)Jejo^*/J | Jackson labs (J. Johnson) | 012882 |
| Ai14 | B6.Cg-Gt(ROSA)26Sor^tm14(CAG-tdTomato)Hze/J^ | Jackson labs (H. Zeng) | 007914 |
| GAD1-EGFP | Not deposited | K. Obata and Y.Yanagawa |  |
| Dlx5/6^FlpE^ | Tg(mI56i-flpe)39Fsh/J | Jackson labs (G. Fishell) | 010815 |
| RCE-FRT | Gt(ROSA)26Sor^tm1.2(CAG-EGFP)Fsh^/Mmjax | Jackson labs (G. Fishell) | 32038 |
| Ai65 | B6;129S-*Gt(ROSA)26Sor^tm65.1(CAG-tdTomato)Hze^*/J | Jackson labs (H. Zeng) | 010815 |
